# Molecular mechanisms of flotillin complexes in organizing membrane microdomains

**DOI:** 10.1101/2024.05.25.595881

**Authors:** Ming-Ao Lu, Yunwen Qian, Liangwen Ma, Jinzhi Hong, Xiaopeng Li, Li Yu, Qiang Guo, Ning Gao

**Author notes:** Correspondence (N.G.), (Q.G.).

## Abstract

Flotillin-1 and flotillin-2 form hetero-oligomers to create flotillin membrane microdomains essential for endocytosis and protein sorting. However, the mechanisms of flotillin oligomerization and microdomain organization remain incompletely understood. Here, we present the cryo-EM structure of human flotillin complex, showing that flotillin-1 and -2 form a 44-mer, membrane attached, and dome-shaped structure that defines a 30-nm circular membrane domain. The cryo-ET data demonstrates that while attached to the cytoplasmic leaflet in situ, flotillin complexes possess intrinsic structural plasticity on the native membrane. Each flotillin complex may represent a fundamental unit of membrane microdomains, with their clustering enabling the formation of larger and more elaborate domains. We further reveal that phosphorylation at residues Y160 (flotillin-1) and Y163 (flotillin-2) may act as a molecular switch to modulate complex assembly, potentially regulating its function in endocytosis. These findings demonstrate the molecular mechanism of flotillin-mediated membrane segregation and microdomain formation, and suggest a previously unrecognized role of flotillin in sequestrating membrane proteins.

## Introduction

Biological membranes are heterogenous in composition and due to the non-uniform distribution of lipids and proteins, are segregated into discrete nanoscale domains that function as dynamic platforms to orchestrate biomolecule interactions both within the membrane and between the membrane and intra/extracellular contents^1,2^. These functional membrane microdomains (FMMs), as basic organizational units of the membrane, are essential for many cellular processes ^3,4^. FMMs are enriched in certain lipid species, such as sterols and sphingolipids and difference classes of proteins^5,6^. Mammalian flotillin-1 and flotillin-2 are two highly homologous proteins found in the membrane microdomains of plasma membrane^7^. They can be efficiently co-immunoprecipitated by each other and extensively colocalize, forming small puncta in the plasma membrane that correspond to flotillin membrane microdomains^8^. The membrane microdomains containing flotillin complexes are laterally mobile within the plasma membrane, and play crucial roles in various cellular processes, including endocytosis^9-11^, signaling^12,13^, protein trafficking^13,14^ and cytoskeleton remodeling^15-17^ (reviewed in^18,19^). The flotillin microdomains can be internalized into the cell with various cargos^8,10,20-22^ or secreted out of the cell^23,24^. For examples, Fyn kinase^11^ and RAB31^23^ respectively regulate the roles of flotillins in endocytosis and exosome secretion. And flotillin overexpression has been implicated in cell invasion and metastasis during tumorigenesis^19,25^. This overexpression disrupts membrane domain homeostasis, promoting plasma membrane invagination and endocytosis towards late endosomes, which in turn alters the trafficking of various cargos^8^, potentially contributing to oncogenic processes.

Although a scaffolding role of flotillins in FMM organization was proposed decades ago^7,26^, the physical basis of flotillin microdomain remains to be an enigma and it is more intriguing that how flotillin is involved in such a variety of different biological processes. Flotillin belongs to the SPFH (Stomatin, Prohibitin, Flotillin, and HflK/C) protein family, which shares a conserved structural organization that comprises an SPFH domain and a subsequent long α-helix (coiled-coil, CC domain) ^27^. These long α-helices of both flotillin-1 and flotillin-2 were demonstrated to be important for their oligomerization^28^. Recent findings suggest that several representative SPFH family proteins can self-oligomerize and anchor to membranes, creating diverse membrane microdomains across various subcellular compartments^27^, such as stomatin complexes in the eukaryotic plasma membrane^29,30^, prohibitin (PHB) complexes in the inner mitochondrial membrane^31-33^, erlin complexes in the endoplasmic reticulum (ER) ^33,34^, and HflK/C complexes in the prokaryotic plasma membrane^35,36^. Different from these typical SPFH proteins, another SPFH protein, bacterial flotillin (also known as FloA) contains a unique membrane-targeting region^37^, and was also demonstrated to play a role in organizing FMMs on bacterial membrane^38,39^. Given the conserved domain architecture of FloA, it likely employs a similar oligomerization mechanism to segregate bacterial membrane^37,40^. While most of these proteins possess transmembrane domains, mammalian flotillins lack such domains yet exhibit robust association with membranes. This association is facilitated by a combination of post translation modifications: S-palmitoylation of a few cysteine residues in their N-terminal domains and an N-terminal myristoylation specific for flotillin-2^15^. In addition, a putative cholesterol recognition/interaction amino acid (CRAC) motif within their SPFH domain may also facilitate membrane association^41^. However, the structural basis for flotillin-1 and flotillin-2 oligomerization and the molecular mechanisms underlying the formation of flotillin membrane microdomains remain unknown.

In this work, we employ multi-modal imaging approaches to resolve the high-resolution structure of purified flotillin complex and the in-situ architecture of the flotillin microdomain. Our results reveal that flotillin complexes exhibit intrinsic structural plasticity on membranes, where each complex may serve as a fundamental unit for microdomain formation. Clustering of these units enables the formation of larger and more elaborate domains. These findings elucidate the molecular basis of flotillin-mediated membrane segregation and uncover their role in membrane protein sequestration, opening avenues for the further exploration of their mechanisms in various cellular functions.

## Results

### Sample preparation and structural determination of the flotillin complex

Flotillin-1 and flotillin-2 were co-expressed in HEK293F cells, and a FLAG tag was placed at the C-terminus of flotillin-1 for the complex purification from solubilized membrane fractions. To further increase sample homogeneity, glycerol density gradient centrifugation was performed and peak fractions were used for structural analysis (Supplementary Fig. 1a). The distribution of flotillin-1 and flotillin-2 in the glycerol fractions was concentrated in high molecular-weight fractions, and each fraction was examined using negative-staining electron microscopy (nsEM) (Supplementary Fig. 1b). The flotillin complex exhibited trapezoid-like shapes (side view) (Supplementary Fig. 1c) and round shapes (top/bottom view) in the nsEM images. From the nsEM images, some complexes were seen to attach to membrane vesicles with the lower bottom of the trapezoids (Supplementary Fig. 1d).

Subsequent cryo-electron microscopy (cryo-EM) analysis confirmed that flotillin complexes resembled a dome-like structure (Supplementary Fig. 2). However, a significant portion of particles exhibited irregularly deformations, and some classes from early 3D classification even show ruptures in the structures (Supplementary Fig. 2d, e). Due to the heterogeneity of the particles, the alignment of 2D particle images during single-particle cryo-EM 3D reconstruction was challenging and resulted in low-resolution density maps. To obtain a high-resolution refinement, we eliminated deformed particles during data processing. Only those particles without apparent deformations were included in the analysis. C22 symmetry was imposed during refinement, which resulted in a final density map of overall 3.57 Å resolution (Supplementary Fig. 2f, 3). Atomic models for flotillin-1 and flotillin-2 were built based on this map.

The structure of the flotillin complex shows that 22 copies of flotillin-1 and 22 copies of flotillin-2 are alternately arranged to form a dome-like structure, denoted as F1_(1)_–F1_(22)_ and F2_(1)_–F2_(22)_, respectively, giving rise to the C22 symmetry (Fig. 1). When viewed from the side, the structure of flotillin complex can be divided into three parts: the base region (∼70 Å high), the wall region (∼140 Å high), and the roof region (∼175 Å wide) (Fig. 1c). The base region comprises the SPFH domains of flotillin-1 and flotillin-2, which are associated with the membrane (density of detergent micelles shown in Fig. 1c). The wall region of the structure is formed by a single layer of parallel-arranged helices, creating a helical barrel. The dome structure is tightly sealed in both the base region and wall region, with the inner leaflet of the plasma membrane serving as the bottom.

**Figure 1.**
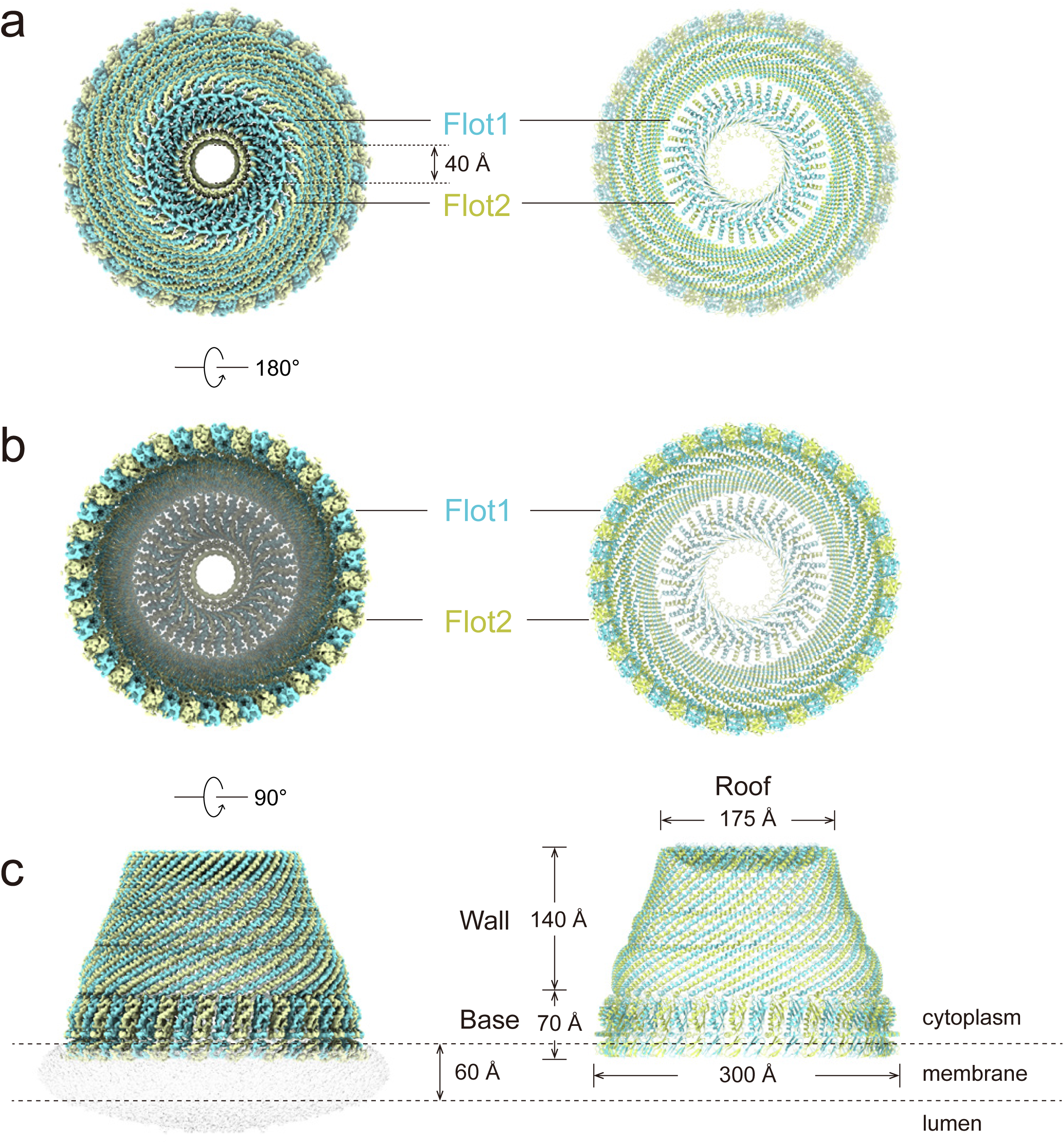
The overall structure of the flotillin complex. **a–c,** Top view (cytoplasmic view) (**a**), bottom view (exoplasmic view) (**b**) and side view (**c**) of the cryo-EM density map (left) and atomic model (right) of the flotillin complex with 22-fold symmetry. The complex is a dome-shaped structure composed of tightly coiled α-helices as the single-layer wall. Subunits of flotillin-1 and flotillin-2 are colored cyan and yellow, respectively.

Recently, concurrent with our work, another study reported the structural analysis of flotillin complexes on isolated membrane vesicles^42^. Their results demonstrate the same overall architecture of the flotillin complex, indicating that the detergent-solubilized flotillin complex has retained its native structure in the membrane environment.

### Structure of the flotillin complex

With approximately 48% sequence identity, flotillin-1 and flotillin-2 have nearly identical 3D structures (Fig. 2a, b). The structures of both proteins can be divided into five domains, including two SPFH domains (SPFH1 and SPFH2), two coiled-coil domains (CC1 and CC2), and a C-terminal domain (CTD), similar to the structure of the HflK/C complex^35,43^.

**Figure 2.**
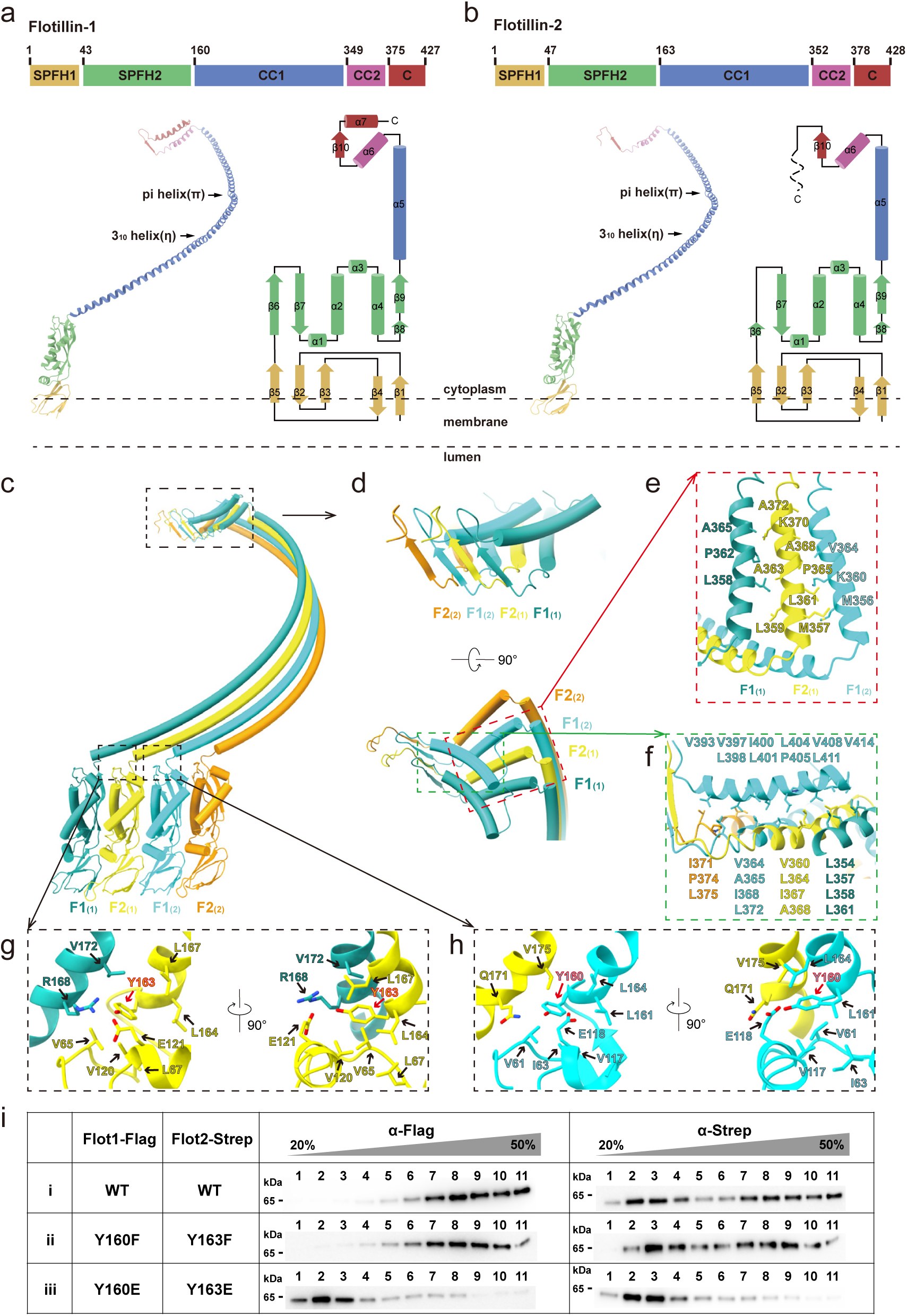
Assembly of the flotillin complex. **a & b,** Schematic illustration of the conserved domain organization of flotillin-1 (**a**) and flotillin-2 (**b**), with individual domains separately colored. SPFH1, gold; SPFH2, green; coiled coil 1 (CC1), blue; CC2, purple; C- terminal domain (CTD), red. **c–h,** Side view (**c**) and zoomed-in views (**d–h**) of the four distinctly colored protomers. **d–f,** Packing of the four protomers in the cap region of the flotillin complex.**e & f,** In the cap region, the CC2 region of F2_(1)_ contacts adjacent CC2 regions of F1_(1)_ and F1_(2)_ (**e**); the α-helix in the CTD region of F1_(2)_ contacts four CC2 regions of F1_(1)_, F2_(1)_, F1_(2)_ and F2_(2)_ (**f**). Residues involved in the detailed interactions are labeled. Hydrophobic interactions are dominant in the cap region. **g & h,** The interfaces between SPFH2 and CC1 domains of adjacent flotillins are stabilized by multiple evolutionarily conserved interactions, including polar contacts that form solvent-exposed surfaces and nonpolar interactions that mediate hydrophobic core packing. Details of these interactions are illustrated for residues surrounding Y163 in F2_(1)_ (**g**) and Y160 in F1_(2)_ (**h**). **i,** Glycerol gradient fractionation analysis of crude membrane extracts co-expressing wild-type (WT) or mutant flotillin constructs (without prior affinity purification; see Methods). Immunoblotting of gradient fractions demonstrates that phosphomimetic mutations (Flot1-Y160E/Flot2-Y163E) completely disrupt dome complex formation. In these experiments, flotillin-1 and flotillin-2 were C-terminally tagged with Flag and Strep epitopes, respectively. Source data are provided as a Source Data file.

Flotillin-1 and flotillin-2 form two distinct heterodimeric interfaces, F1:F2 and F2:F1 (Fig. 2c). The SPFH1 domains of flotillin-1 and flotillin-2 interact with each other via the less conserved loop regions on their structurally conserved five-stranded core (Supplementary Figs. 4a–d and 5). The interfaces of adjacent SPFH2 domains are characterized by oppositely charged surface patches, which exhibit a clear charge complementarity (Supplementary Fig. 4e–h). These charged patch pairs dictate that flotillin-1 and flotillin-2 should be alternately arranged to form a complete complex. Following the SPFH domains, the CC1 domains of flotillin-1 and flotillin-2 both contain 189 residues, which are significantly longer than those of other SPFH family members (Supplementary Fig. 5), and they align perfectly to form a right-handed helical barrel and serve as the dome wall (Fig. 1). The interactions between parallel CC1 domains are evenly distributed along the entire helices, with no preference for either polar (Supplementary Fig. 6a–c) or hydrophobic interactions (Supplementary Fig. 6d–f). Besides, there are two small bends at the sites of Pi-helical (π) and 3_10_-helical (η) motifs of CC1 domains (Fig. 2a, b), constituting the two hoops observed in the side view of the dome (Fig. 1c).

Following the CC1 domains, the CC2 domains and CTDs of both flotillins form the roof of the dome, primarily stabilized by hydrophobic interactions (Fig. 2d–f). The 44 CC2 domains (α6) converge towards the center, where 44 β-strands assemble into a β-barrel core. The subsequent sequences in the CTDs of flotillin-1 and flotillin-2 exhibit distinct structural characteristics, resembling the organization observed in the HflK/C complex^35,43^. In flotillin-1, the helices (α7) in the CTDs are oriented outwards from the center, with the C-terminus exposed to the cytoplasmic space. In contrast, the putative helices (α7) in the CTDs of flotillin-2 appear to be buried within the flotillin complex, although their exact conformation is not fully resolved in the map. This structural difference aligns with experimental observations, as attempts to purify the flotillin complex using a FLAG tag at the C-terminus of flotillin-2 were unsuccessful, suggesting that the C-terminal region of flotillin-2 is inaccessible in the assembled complex. Similar to the HflK/C complex^43^, progressive truncation of the C-terminal sequences may disrupts dome structure formation (Supplementary Fig. 7).

### Potential phosphorylation mediated regulation on the flotillin complex assembly

Y160 in flotillin-1 and Y163 in flotillin-2 are two known phosphorylation sites for Src family kinase Fyn^6^. These two residues are localized in the hinge regions between the SPFH2 and CC1 domains of flotillin-1 and -2, which might play a role in determining the orientation of the CC1 domain relative to the SPFH2 domain. Structure-wise, Y163 and Y160 are stabilized by multiple conserved hydrophilic and hydrophobic interactions across both flotillin homologs, in the F1_(1)_:F2_(1)_ and F2_(1)_:F1_(2)_ interfaces, respectively (Fig. 2g, h). On the one hand, polar interactions are established between the hydroxyl group of Y163 of F2_(1)_ and R168 of F1_(1)_ (Fig. 2g), and between the hydroxyl group of Y160 of F1_(2)_ and Q171 of F2_(1)_ (Fig. 2h), forming the solvent-exposed surfaces. On the other hand, the benzene rings Y163 and Y160 directly contribute to the formation of a hydrophobic core involving both inter- and intra-molecular interactions, including V172 of F1_(1)_ with V65, L67, V120, L164, and L167 of F2_(1)_ (Fig. 2g), and V175 of F1_(2)_ with V61, I63, V117, L161, and L164 of F2_(1)_ (Fig. 2h), respectively.

Next, we introduced mutations on Y160 of flotillin-1 and Y163 of flotillin-2 and examined their effects on the complex assembly. Since there is no phosphomimetic mutations for tyrosine, we introduced Y160E and Y163E to partially mimic the charge effect of phosphorylation, and found that they completely abolished the complex assembly (Fig. 2i, Supplementary Fig. 7). In contrast, neither the tyrosine mutations (Y160F of flotillin-1 and Y163F of flotillin-2) nor the double mutations (Y160F/R168A of flotillin-1 and Y163F/Q171A of flotillin-2) affected flotillin complex assembly (Fig. 2i, Supplementary Fig. 7). These results indicate that the stability of the Y160/Y163 participated hydrophobic core is essential for the formation of the flotillin complex. Very interestingly, it was previously shown that Y160F (flotillin-1) and Y163F (flotillin-2) mutations could block Fyn-regulated flotillin-mediated endocytosis^11,17^. In combination with our results, it appears to suggest that flotillin-mediated endocytosis might involve the disassembly of the dome structure.

Altogether, the previous and our observations indicate that phosphorylation at these tyrosines may destabilize the local hydrophobic cores, inducing a change on the orientation between the SPFH2 and CC1 domains that could function as a molecular switch to trigger the flotillin complex disassembly.

### Flotillin complex mediated membrane compartmentalization

The SPFH1 domains mediate the membrane association of flotillins, with approximately half embedded in the detergent micelle (Fig. 3a–d). These domains are rich in aromatic residues, including F2, F3, F16 and F30 of flotillin-1, and W32, W34, W36 and W37 of flotillin-2 (Fig. 3d), contributing to the highly hydrophobic nature of the bottom rim of the flotillin complex (Fig. 3e). Notably, palmitoylated cystine residues (C34 of flotillin-1; C4, C19 and C20 of flotillin-2)^15^ and myristoylated glycine residue (G2) of flotillin-2 are all buried in membrane (Fig. 3d). Together, these features collectively ensure the stable and efficient membrane association of flotillins.

**Figure 3.**
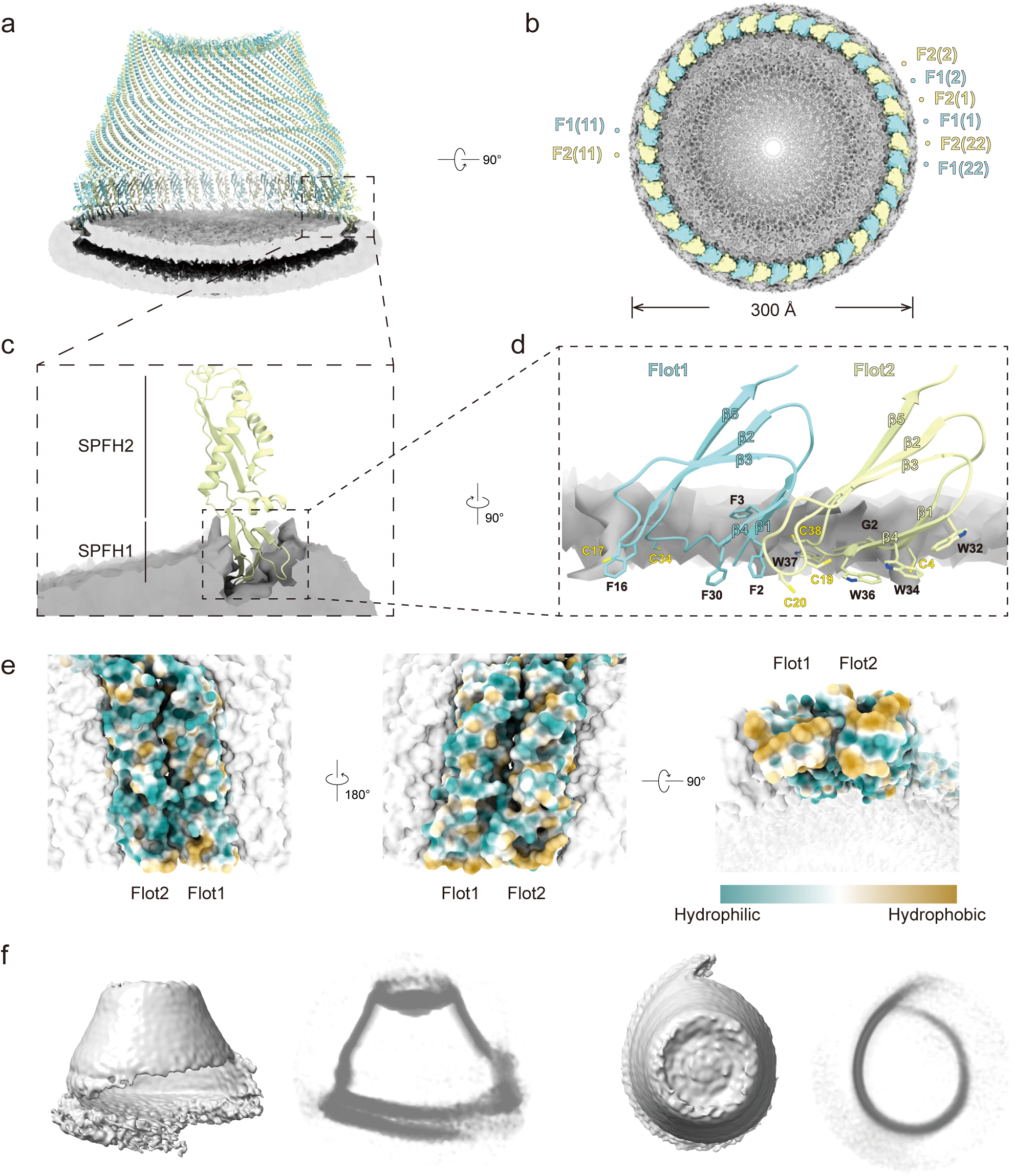
Flotillin complex mediated membrane compartmentalization. **a–d,** SPFH1 domains of flotillin-1 and flotillin-2 are inserted in the cytoplasmic leaflet of the membrane. Flotillin-1 and flotillin-2 are colored cyan and yellow, respectively. Detergent density is colored gray. **a,** Vertical cross-section view of the atomic model of the flotillin complex with the low-pass filtered cryo-EM density map of detergent micelle superimposed, in the position of the axis of symmetry. **b,** Horizontal cross-section view of the low-pass filtered cryo-EM density map of the flotillin complex with detergent micelle, in the position of the SPFH1 region. Protomers of flotillin-1 and flotillin-2 are numbered anti-clockwisely. **c,** Zoomed-in view of the SPFH regions of the flotillin complex. **d,** Zoomed-in view of the SPFH1 regions of the flotillin complex with ninety-degree rotation, with the removal of detergent density in the foreground. Aromatic residues (F2, F3, F16 and F30 of flotillin-1; W32, W34, W36 and W37 of flotillin-2) are inserted in the membrane. Cystine residues (C34 of flotillin-1; C4, C19 and C20 of flotillin-2), which are reported to be palmitoylated, and glycine residue (G2 of flotillin-2), which is reported to be myristoylated, are all buried in membrane. **e,** Hydrophobic surface model of the SPFH regions from a flotillin-1/2 dimer within the complex. The PTMs in this region are not considered here. Cyan, hydrophilic; white, neutral; gold, hydrophobic. **f,** Side views and top views of deformed maps during cryo-EM SPA 3D classification. The slabs of these maps are shown aside, as illustrated in Supplementary Fig. 2d, e.

The 44 circularly arranged SPFH1 domains define a 30 nm-diameter membrane area on the cytosolic leaflet of the membrane (Fig. 3a, b). When all flotillin subunits are precisely aligned, they form a sealed dome that effectively compartmentalizes both the membrane and the perimembrane space, physically isolating it from the surrounding environment (Fig. 1). However, in addition to these well-aligned complexes, irregular deformations and structural ruptures were frequently observed during in vitro structural determination (Fig. 3f, Supplementary Fig. 2d–f). These observations suggest that the sealed dome may dynamically open and reseal, potentially regulating access to the segregated region.

### In situ structure of the flotillin complex

To explore whether such deformations occur within a cellular context and to gain further insight into the potential functional roles of the flotillin complex, we developed a stable Jurkat T cell line co-expressing F1-EGFP and F2-mCherry for in situ studies. Flotillin complexes are known to exhibit a polarized distribution in T cells, localizing to caps, uropods, and the immune synapse during resting^44^, migrating^16^, and conjugated stages^13^, respectively. Consistently, our fluorescent confocal imaging results show that flotillin-1 and flotillin-2 were co-localized in caps or uropods of Jurkat T cells only when both flotillin homologs are co-expressed (Fig. 4a, Supplementary Fig. 8a–c), highlighting the necessity of heterooligomer formation for their function. Additionally, glycerol density gradient centrifugation assay confirmed that the fluorescent tags did not disrupt the assembly of flotillin complexes (Supplementary Fig. 8d).

**Figure 4.**
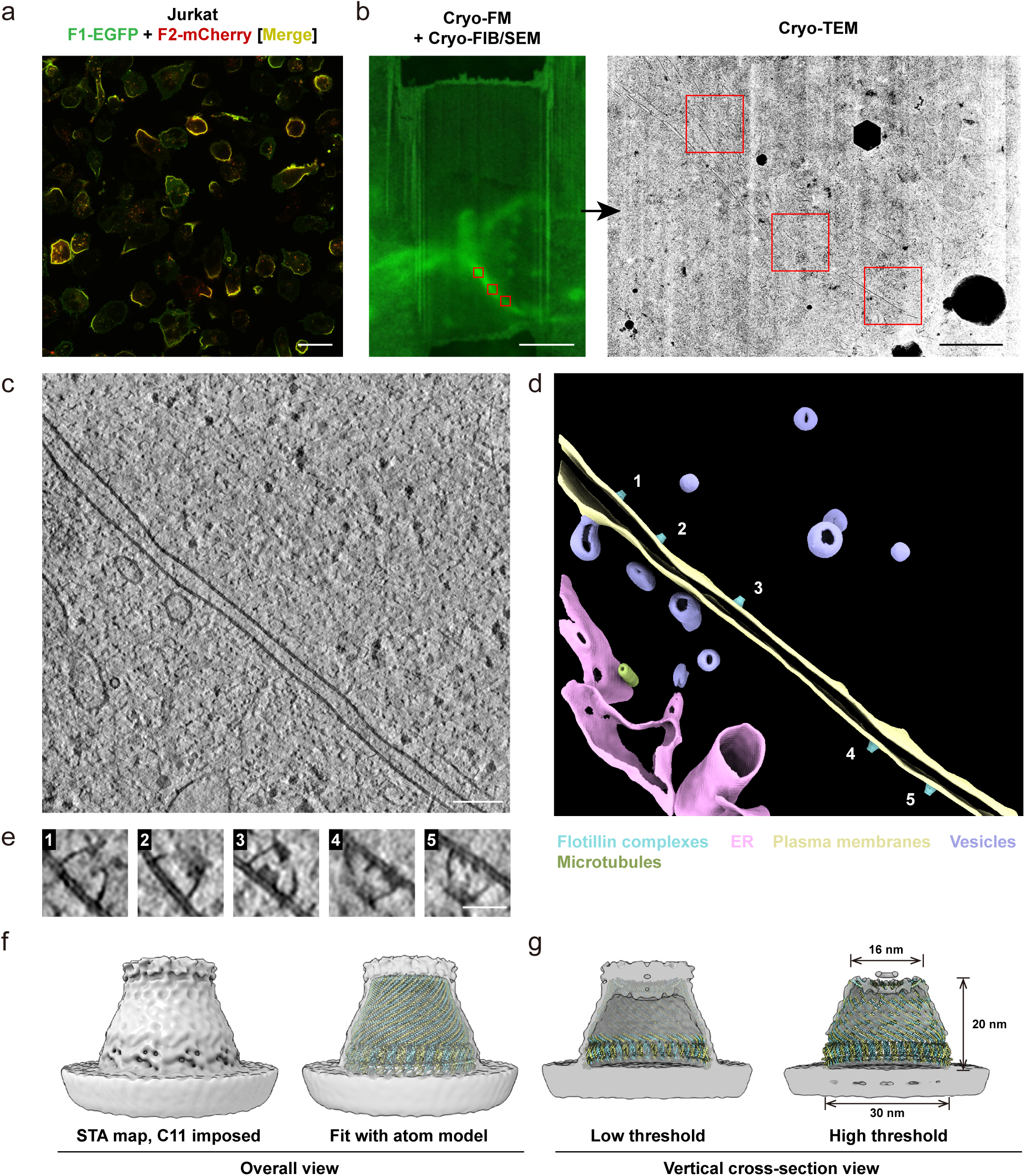
Visualization of the flotillin complex in situ. **a,** Jurkat cells stably expressing flotillin-1-EGFP and flotillin-2-mCherry exhibit colocalization at the caps or uropods, as visualized by fluorescence microcopy. **b,** Correlative cryo-light and electron microscopy workflow. Regions of interest identified by fluorescence microscopy serve as guide for subsequent tomographic tilt series collection by cryo-EM. **c,** A 1.2 nm thick tomographic slice acquired within the area containing the GFP signal (corresponding to flotillin complexes). **d,** 3D rendering of the tomogram shown in **c**. Flotillin complexes, plasma membranes, vesicles, ER and microtubules are colored as indicated. Selected flotillin complexes magnified in **e** are indicated by numbers. See **Supplementary Movie 1** for animated visualization. **e,** Tomographic slices zoomed in on the representative flotillin complexes identified within the tomogram shown in panel (**c**). Notably, the side views of these complexes exhibit variability in shape. Additionally, some complexes (3 and 4, for example) contain internal regions of increased electron density. **f & g,** Subtomogram averaging result of flotillin complexes (**f**) and its vertical cross-section views (**g**) at different contour levels. The model of flotillin complexes built with SPA map was fitted in to the STA maps for comparation. Scale bars, 20 µm (**a**), 5 µm (left panel in **b**), 1 µm (right panel in **b**), 120 nm (**c**) and 30 nm (**e**).

Guided by the fluorescent signals of F1-EGFP and F2-mCherry, lamellae were prepared at regions enriched with flotillin complexes (Fig. 4b). Subsequent tomographic reconstructions revealed a cellular periphery with a clearly visible plasma membrane, consistent with the known localization of flotillin complexes in caps or uropods (Fig. 4c, d, Supplementary Movie 1). Notably, a significant number of hollow dome-shaped structures with heights of approximately 20 nm were observed attached to the cytosolic side of the plasma membrane (Fig. 4e), resembling our cryo-EM structure of the flotillin complex. To validate the identity of these structures, sub-tomogram averaging was performed. While the limited data size restricts the final map resolution and precludes definitive identification based solely on structure, the overall shape of the structure in conjunction with the correlative light and electron microscopy (CLEM) results strongly suggest their formation by the flotillin complex (Fig. 4f, g).

Individual subtomograms revealed additional electron densities within the complexes (Fig. 4e). These densities likely correspond to various partner proteins. However, they were absent in the averaged structure, suggesting a high heterogeneity of partner proteins with the domes. Additionally, a significant degree of structural variation was observed among the individual complexes (Supplementary Fig. 9a–b, 10a). This observation aligns with the findings from purified samples (Fig. 3f, Supplementary Fig. 2b–e, 9c–d), further supporting the notion that the flotillin complexes are highly dynamic in nature. This property might be essential for the flotillin complex’s cellular functions.

### Organization model of flotillin membrane microdomains

The tomographic data offered a unique opportunity to investigate the spatial distribution and membrane preferences of flotillin complexes in situ. A majority of the observed flotillin complexes existed as individual units. Given the substantial membrane area encompassed by the dome-shaped complex, it is conceivable that the complex, along with its associated proteins, contribute to the organization of a specialized membrane microdomain. Notably, we also observed clusters of flotillin complexes on extracellular vesicles (Fig. 5a, Supplementary Movie 2), endosomes and plasma membrane (Fig. 5b, Supplementary Movie 3), which aligns with the presence of larger flotillin membrane microdomains implicated in exosome formation^45^ and endocytosis^8,10,11^.

**Figure 5.**
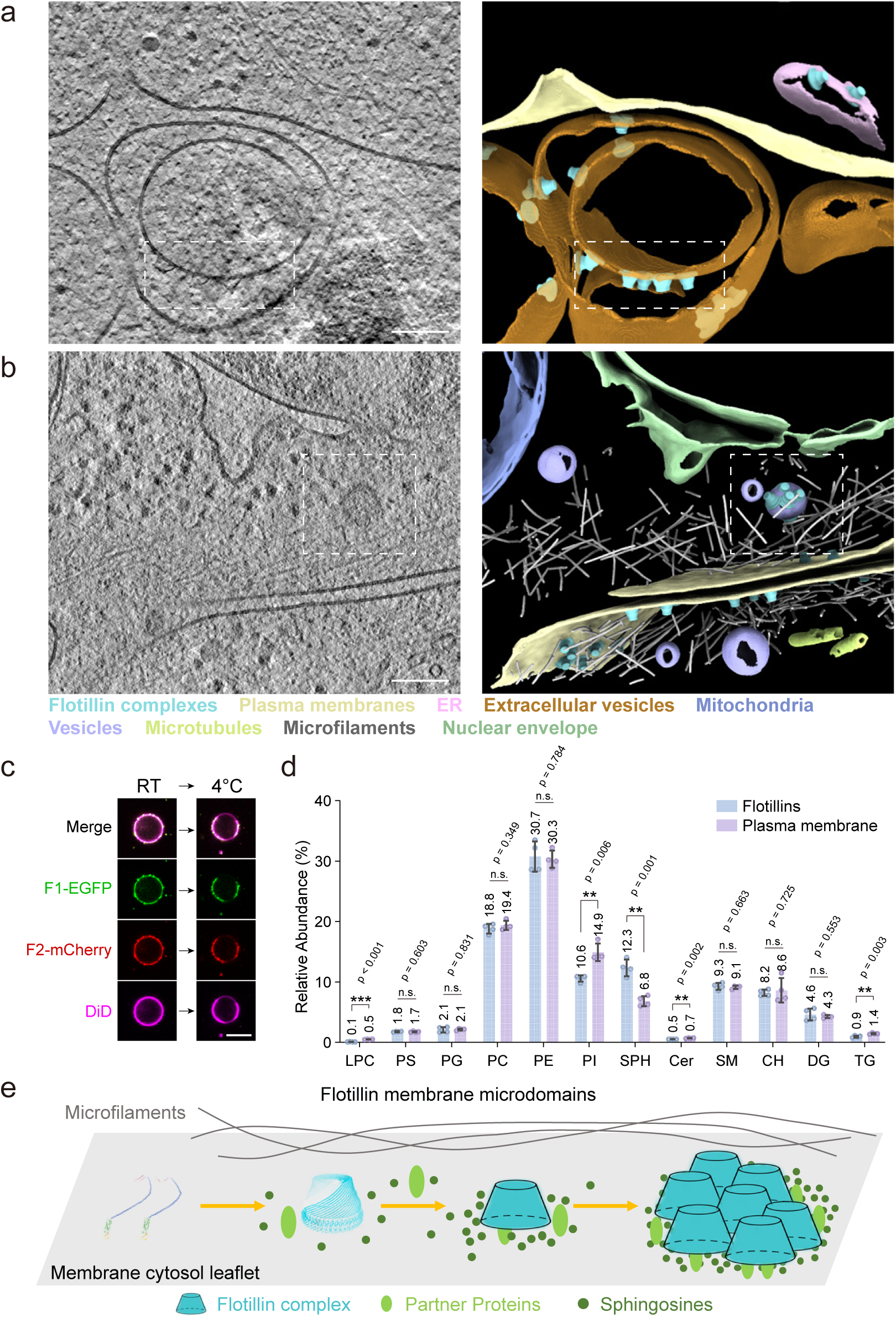
Organization of membrane microdomains by flotillin complexes. **a & b,** Tomographic slice (left) and corresponding 3D reconstruction (right) from representative tomograms of regions marked by flotillin complex fluorescence. Flotillin complexes localize to the cytosolic leaflet of the plasma membrane and endosomes, as well as the luminal leaflet of extracellular vesicles (**a**), and are also observed in close association with actin filaments (**b**). The boxed areas highlight clusters of flotillin complexes within extracellular vesicles (**a**) and endosomes (**b**). Animated views of (**a**) and (**b**) are provided in **Supplementary Movies 2** and **3**, respectively. Subcellular components are colored as indicated. **c,** After NEM treatment, GPMVs were isolated from cells stably expressing F1-EGFP and F2-mCherry, and separated into coexisting Lo/Ld phases as the temperature was lowered from room temperature to 4°C. DiD dye marks Ld phase. **d**, Comparative lipidomics analysis of endogenous lipids in purified flotillin samples and the plasma membrane fraction. Data are presented as mean ± SD from two biological replicates with two technical replicates each (individual data points represented as dots). Two-sided unpaired t-tests with Welch’s correction were performed without multiple comparisons correction. ** *p* < 0.01, \*\*\**p* < 0.001, n.s.: not statistically significant. Complete statistical information for null hypothesis testing is provided in **Supplementary Data 1**. LPC, lysophosphatidic acid; PS, phosphatidylserine; PG, phosphatidylglycerol; PC, phosphatidylcholine; PE, phosphatidylethanolamine; PI, phosphatidylinositol; SPH, sphingosine; Cer, ceramide; SM, sphingomyelin; CH, cholesterol; DG, diacylglycerol; TG, triacylglycerol. **e,** Proposed model of flotillin membrane microdomain. Each minimal microdomain unit consists of a flotillin complex together with recruited partner proteins and sphingolipids. The unit can cluster with others, driving lateral segregation within the membrane. Through this progressive clustering, larger functional membrane microdomains are ultimately formed. Tomographic slices are 1.2 nm thick. Scale bars, 120 nm (**a** and **b**), 10 µm (**c**)

The flotillin complexes were observed on membranes with varying curvature in situ. Most flotillin particles were located on relatively flat membranes (Supplementary Fig. 10a), and some appeared on convex or concave vesicle surfaces (Supplementary Fig. 10b, c). In purified samples, the averaged structure of non-vesicle-associated particles retained a concave-shaped membrane at the bottom (Supplementary Fig. 10d), while some complexes attached to the convex membrane surfaces of vesicles (Supplementary Fig. 10e). These observations indicate that flotillin complexes exhibit no specific preference for membrane curvature.

Since flotillin complexes also exhibit a clustering pattern, we sought to examine whether this behavior is related to the distribution of certain lipid species on the plasma membrane. Notably, flotillin was considered as a marker of lipid rafts^7,26,27^, which commonly refer to those highly dynamic, cholesterol and sphingolipids enriched microdomains (ranging from 10 to 200 nm) on the plasma membrane^46^. A model system to study the lipid driven phase separation property of plasma membrane is the giant plasma membrane vesicle (GPMV)^47^. Therefore, we prepared GPMVs from human Jurkat cells stably expressing tagged flotillin-1 and flotillin-2. Previously, flotillins were found to be enriched in the detergent-resistant membrane (DRM) fractions^7,26^, a sign usually suggesting a localization in the liquid-ordered (Lo) phase of plasma membrane. Unexpectedly, but consistent with a previous report on flotillin-2 ^48^, our data indicate that both flotillin-1 and -2 predominantly partition into the liquid-disordered (Ld) phase in GPMV-based partitioning assay (Fig. 5c, Supplementary Fig. 11a). Similarly, tetraspanin-4 (T4), a member of the tetraspanin protein family that scaffolds tetraspanin-enriched microdomains^49,50^, colocalizes with flotillins in the vesicles within L929 cells (Supplementary Fig. 11b), and also partitions into the Ld phase (Supplementary Fig. 11c).

Furthermore, the use of GDN detergents during protein purification preserved portions of the membrane within the complex (Supplementary Fig. 1c), enabling mass spectrometry to identify endogenous lipids co-purified with flotillins (Supplementary Data 1). Comparative lipidomics analysis revealed a 1.8-fold enrichment of sphingosine (SPH) in the purified flotillin sample compared to the plasma membrane fraction, with no significant increase observed in other raft-associated lipids, such as cholesterol (CH) or other sphingolipids. This enrichment was accompanied by relative decreases in several other lipid species, including phosphatidylinositol (PI), lysophosphatidylcholine (LPC), ceramide (Cer), and triacylglycerol (TG) (Fig. 5d). This observation aligns with previous reports indicating that flotillins interact with sphingosine^51^. These findings, together with results of GPMV partitioning assay suggest that the flotillin complex does not have a strong preference for certain lipid-enriched membrane regions and its further clustering may not primarily dependent on raft-lipid mediated segregation mechanism^3^.

Here, we propose a model of the flotillin membrane microdomain (Fig. 5e): The membrane encompassed by the flotillin complex is laterally segregated as a basic unit of the microdomain (30 nm in diameter). Together with specifically associated sphingosine and proteins, the flotillin membrane microdomain could exert diverse functions in different cellular processes. Under certain conditions, or dependent on contextual factors, flotillin complexes could cluster to form larger membrane microdomains (over 200 nm in diameter), serving as a platform for signaling or inter-membrane trafficking.

## Discussion

Our cryo-EM analysis reveals the structure of the flotillin complex obtained from detergent-solubilized, affinity-purified samples. In parallel, Fu & MacKinnon employed an innovative approach, isolating flotillin complexes directly from cell-derived vesicles while maintaining a near-native membrane environment without detergent disruption^42^. In terms of the structure, the two studies are highly consistent, indicating that the oligomerization and subunit stoichiometry are intrinsic properties of flotillin proteins. Together, the two complementary studies demonstrates that the flotillin complex acts as a physical container that compartmentalizes the membrane and perimembrane space. Structurally, it forms a dome-shaped architecture, with SPFH1, SPFH2, and CC1 domains composing the walls, the cytoplasmic leaflet of the lipid bilayer serving as the base, and CC2 and C-terminal domains forming the roof. A 4-nm hole observed at the center of the roof is likely filled by the flexible CTDs of flotillin-2. The sealed dome structure creates a physically isolated membrane compartment that provides a confined microenvironment for specialized molecular processes. The observed densities within the dome suggest potential encapsulation of partner proteins, analogous to the FtsH proteins in HflK/C complexes^43^, aquaporin-1 and urea transporter-B in Stomatin complexes^30^ and *m*-AAA protease in PHB complexes^32^. This encapsulation likely requires dynamic opening and resealing of the dome structure, consistent with its demonstrated flexibility and deformability in both in vitro and in situ observations. Similar structural behavior has been reported for other SPFH family complexes, including HflK/C^43,52^, vault^53^, prohibitin-1/2^31-33^, stomatin^29,30^ and erlin-1/2^33,34^ complexes, highlighting the functional importance of dynamic structural adaptability. Furthermore, phosphorylation at Y160 (flotillin-1) and Y163 (flotillin-2) may disrupt the hydrophobic core-mediated SPFH-CC1 domain orientation, potentially serving as a phosphorylation-dependent regulatory mechanism for complex disassembly, which could play a crucial role in Fyn kinase-regulated flotillin-mediated endocytosis^11,17^.

Interestingly, contrary to the traditional view of flotillins as lipid raft markers^7,26,27^, flotillins partition into the non-raft phase in GPMVs and show limited association with typical raft-associated lipids such as cholesterol and sphingomyelin, except for sphingosines. It is well accepted that lipid raft formation is driven primarily by lipid-lipid interactions^6^. Flotillin complexes, instead of directly organizing lipid domains, appear to function as structural elements that enable lateral compartmentalization of the membrane. Their raft-phase localization or clustering in vivo likely depends on contextual factors, including local lipid composition and their interactions with other membrane or non-membrane components, such as the cytoskeleton^18,54^.

The enrichment of sphingosines within flotillin microdomains offers a potential extension of their role. Flotillins may stabilize sphingosines within these microdomains^51^, providing a platform for biochemical processes such as the recruitment of sphingosine kinase 2 and subsequent production of sphingosine-1-phosphate (S1P)^55^. This process potentially links flotillin-mediated functions, including endocytosis^56^ and gene expression regulation^51^, to the modulation of S1P-dependent signaling pathways^18^. However, this connection remains speculative and requires further investigation to fully elucidate its biological relevance.

Compared with the recent work by Fu et al.^42^, our study has also revealed the intrinsic plasticity and membrane clustering behavior of flotillin complexes via in situ cryo-electron tomography. There are a few interesting differences between two studies. First, their study suggests that flotillin complexes interact with membranes via both the N-terminal basal region and the C-terminal roof region^42^. In contrast, our analysis of over 200 flotillin complexes in situ found no evidence of roof region interacting with the membrane. This difference can be rationalized by different conditions of the two studies. The roof region of the flotillin complex features a concave surface densely enriched with negative charges. In their study, 2 mM Ca²⁺ used in GPMV preparation—approximately 200-fold higher than physiological levels^57^—would neutralize the negative charges on both the phospholipid membranes and the roof region, and could potentially mediate the interaction between the membrane and the roof. In addition, the high vesicle concentration used for cryo-EM sample preparation might further promote their interaction. Second, their study proposes that flotillin complexes induce membrane curvature through conformational changes^42^. Drawing on structural plasticity of flotillin complexes observed both in vitro and in situ, we believe that this behavior reflects an adaptation to existing membrane curvatures rather than an intrinsic ability to induce curvature. These different interpretations highlight importance of in situ studies in capturing the native behavior of flotillin complexes.

In summary, flotillin complexes act as structural units that compartmentalize membranes, creating confined and dynamic environments for specific molecular interactions with potential implications for cellular signaling. These findings broaden our understanding of SPFH family proteins, highlighting their common roles in membrane microdomain organization and membrane protein compartmentalization, and provide a basis to further dissect regulatory mechanisms of flotillins in diverse cellular functions.

## Methods

### Cloning, expression and purification of the flotillin complex

The coding DNA sequences for human flotillin-1 (UniProt ID: O75955) and flotillin-2 (UniProt ID: Q14254) were individually cloned into the pMLink vectors at the multiple cloning sites. Flotillin-1 contains a C-terminal FLAG tag, and flotillin-2 has no tag or a C-terminal Strep tag. Using the LINK sequences in pMLink vectors^58^, after digestion with PacI and SwaI separately, the two genes in different pMLink vectors were cloned into the same pMLink vector via ligation-independent cloning (LIC) method^59,60^. To co-express both flotillins, the pMLink-flotillin-1 and pMLink-flotillin-2 vectors were combined in this way to generate the pMLink-flotillin-1/2 vector. Mutations and truncations of these genes were introduced individually into the pMLink-flotillin-1 and pMLink-flotillin-2 vectors prior to the combination.

HEK 293F cells were cultured in SMM 293T-II medium (Sino Biological Inc.) at 37°C under 5% CO_2_. When the cell density reached 1.5 × 10^6^ cell/mL, the cells were transfected with the pMLink plasmid containing two flotillins. For one liter of cell culture, 1 mg of plasmid and 4 mg of polyethylenimine (Polysciences) were preincubated in 50 ml of fresh SMM 293-TII medium for 30 min. The mixture was then added to the cell culture. After 48 hours of incubation, 6 liters of the cell were harvested and resuspended in the prechilled Buffer A (40 mM Tris-HCl pH 7.4, 150 mM NaCl, 0.5 mM Mg(OAc)_2_, 10% glycerol) supplemented with 1% (v/v) protease inhibitor cocktail (Mei5bio). After sonication, the sample was subjected to centrifugation at 3,000 × g for 30 min to remove unbroken cells and large debris. Membranes were then pelleted by ultra-centrifugation at 50,000× rpm for 1 h (rotor 70 Ti, Beckman Coulter). All pelleted membranes were solubilized with Buffer B (Buffer A supplemented with 1% (w/v) glyco-diosgenin (GDN, Anatrace)) for 2 h. After centrifugation at 8,000 × g for 10 min, the supernatant was incubated with anti-FLAG M2 affinity resin (Sigma) at 4°C for 2 h. After rinsing the resin with wash Buffer C (Buffer A supplemented with 0.004% (w/v) GDN), the protein was eluted with Buffer D (Buffer C supplemented with 0.5 mg/ml 3× FLAG peptide) and concentrated to ∼100 μL using a 100-kDa-cutoff spin concentrator (Millipore).

Then, the samples were loaded on a 20%–50% glycerol density gradient and subjected to ultra-centrifugation at 100,000× g for 14 h (TLS-55 rotor, Beckman Coulter). All fractions of the glycerol gradient were collected manually in separate tubes, with 200 μL per fraction and analyzed with 10% SDS-polyacrylamide gel electrophoresis (SDS-PAGE). Fractions with highest homogeneity, as verified by negative staining electron microscopy, were pooled and dialyzed into Buffer E (40 mM Tris-HCl pH 7.4, 150 mM NaCl, 0.5 mM Mg(OAc)_2_, 5% glycerol, 0.004% (w/v) GDN). At last, the purified protein samples were concentrated to a desired concentration using a 100-kDa-cutoff spin concentrator (Millipore), and then stored at −80°C.

### Glycerol gradient fractionation analysis of crude membrane extracts

To evaluate whether the designed mutations disrupt dome structure formation, we adapted the aforementioned protocol while omitting the affinity purification step. Briefly, HEK293F cells expressing wild-type (WT) or mutant flotillin constructs (individually or in combination) were harvested and resuspended in Buffer A supplemented with 1% (v/v) protease inhibitor cocktail (Mei5bio). Cell lysis was performed using a French press, followed by clarification centrifugation at 3,000 × g for 30 min. Membrane fractions were subsequently pelleted by ultracentrifugation at 45,000 rpm (70 Ti rotor, Beckman Coulter) for 1 h at 4°C. The membrane pellets were solubilized overnight at 4°C in Buffer F (Buffer A containing 1% (w/v) n-dodecyl-ß-D-maltoside (DDM, Anatrace)), followed by clearance centrifugation at 52,000 rpm (TLA-55 rotor, Beckman Coulter) for 25 min. The resulting supernatant was layered onto a 20%-50% glycerol density gradient and subjected to ultra-centrifugation at 52,000 rpm (TLS-55 rotor, Beckman Coulter) for 5 h at 4°C. Gradient fractions (200 μL each) were manually collected and analyzed by SDS-PAGE and western blotting using anti-Flag antibody (Sigma-Aldrich, F3165) for flotillin-1-Flag and anti-Strep antibody (GenScript, A01732) for flotillin-2-Strep.

### Stable cell line construction, GPMV preparation and microscopy imaging

For lentivirus packaging, HEK293T cells were plated in 10 cm dishes (4 × 10^6^ cell/dish) and cultured in DMEM (Gibco), penicillin-streptomycin, and 10% fetal bovine serum (Sigma). A transfection mix was set up as follows: 10 µg transfer vector (pLVX-F1-EGFP, pLVX-F2-mCherry or pLVX-T4-BFP), 15 µg psPAX2 and 5 µg pVSVG in 0.5 mL Opti-MEM were combined with 60 µg PEI in 0.5 mL Opti-MEM (Thermo Fisher) and incubated for 20 minutes. Cell media was removed, and 9 mL Opti-MEM and 1 mL of transfection mix were added per dish. The transfection media was replaced after 6 h by plating media supplemented with 10% fetal bovine serum, and the supernatant was collected after additional 48 h. 10 mL lentivirus containing supernatant was concentrated to 1 mL using a 100-kDa-cutoff spin concentrator (Millipore) at 1,500 × g and freshly used for infection after being supplemented with equal volume of complete medium.

Jurkat cells were suspended cultured in RPMI 1640 (Gibco), penicillin-streptomycin, and 10% fetal bovine serum (Sigma) at 37°C under 5% CO_2_. When the cell density reached 0.25 × 10^6^ cell/mL, the cells were infected with F1-EGFP and/or F2-mCherry lentivirus. Infected cells were sorted by Astrios EQ (Beckman Coulter) based on the corresponding fluorescence and cultured as normal. L929 cells stably expressing T4-BFP, F1-EGFP and F2-mCherry were also constructed in this way. To adhere Jurkat cells to dishes or cryo-EM grids, 50 µg/ml poly-D-lysine (Thermo Fisher) or 10 µg/ml fibronectin (Yeasen) solutions were added to the culture surface. After incubating 1 h at 37°C, the coating reagents were removed and the dishes were freshly used for Jurkat cell adhering.

GPMVs were isolated and imaged as described previously ^47,61^. Briefly, cell membranes of Jurkat or L929 stable cell lines were labeled with 5 µg/ml fluorescent liquid-disordered phase or liquid-ordered phase marker for 20 min at 37°C. Then, cells were washed in GPMV buffer (10 mM HEPES, 150 mM NaCl, 2 mM CaCl_2_, pH 7.4) and incubated with GPMV buffer supplemented with 25 mM paraformaldehyde (PFA) and 2 mM dithiothreitol (DTT) (or 2 mM N-ethylmaleimide (NEM) alone) for 1 h at 37°C. The suspension was centrifuged at 100× g for 1 min, to pellet cell debris. The GPMV supernatant was transferred into a microcentrifuge tube by pipetting, and 400 µM deoxycholic acid (DCA) was added to increase phase separation temperature of GPMV membranes^62,63^. The GPMV suspension was left in the microcentrifuge tube for 20–30 min, and GPMVs would concentrate at the bottom of the tube. The sample was pipetted directly from the bottom of the tube for further microscopy experiments.

For confocal microscopy, cells or GPMVs were cultured or loaded in 35 mm Glass bottom dish (Cellvis), and imaged using Nikon A1RSi+ laser scanning microscope with 60× and 100× objective under oil and lasers exciting at 405 nm, 488 nm, 561 nm and 640 nm. To visualize polarized uropod or “cap” structure of Jurkat cells, fibronectin-coated dishes were used to promote cell adhesion. Polarization was induced by treatment with 100 ng/mL RANTES (MedChemExpress) for 4 h at 37°C/5% CO₂ to stimulate migratory morphology. Nuclei were counterstained with Hoechst 33342 prior to live-cell imaging. To view proteins and lipids partitioning on GPMV membranes, the temperature was decreased below 15°C for GPMVs derived with PFA/DTT and below 5°C for the ones derived with NEM. Image analysis was conducted using NIS-Elements Viewer (Nikon) and ImageJ ^64^.

### Cryo-EM single particle sample preparation, image collection

The sample homogeneity was examined by negative staining electron microscopy. Four microliters of samples were applied onto copper grids and stained with 2% uranyl acetate. The grids were examined with an FEI Tecnai T20 TEM at 120 kV.

Cryo-EM grids were prepared with GraFuture^TM^ RGO TEM Grid (GraFuture-RGO001A, Shuimu BioSciences Ltd., CN). The grids were glow-discharged for 10 s with a plasma cleaner prior to sample freezing. Three microliters of freshly prepared flotillin complex samples (∼0.1 mg/mL) were applied onto the grids mounted in the chamber of an Thermo Fisher Vitrobot Mark IV at 16°C with 100% humidity (adsorption time 1 s, waiting time 60 s, and force -1), and flash-frozen in liquid ethane cooled by liquid nitrogen. Grids were screened on Thermo Fisher Talos Arctica (Gatan K2 Summit detector), operated at 200 kV. Images were collected with a 300 kV Thermo Fisher Titan Krios TEM (Gatan K3 summit detector, with GIF energy filter). The movies were acquired using EPU (Thermo Fisher Scientific) at a dose rate of 8.52 e^-^/s/Å^2^ and an exposure time of 4.81 s with 32 frames. The data was collected at the magnification of 64,000× corresponding to a physical pixel size of 1.37 Å/pixel and with defocus ranging from -1.6 to -2.2 μm. Two datasets of movie stacks (19,477 in total) were collected separately.

### Cryo-EM single particle analysis (SPA)

The motion correction and electron-dose weighting of the micrographs were performed with MotionCor2^65^ and the contrast transfer function (CTF) parameters were estimated with the program of Gctf^66^. Particle picking, classification and structure refinement were done with RELION4.0^67^.

The two datasets underwent separate processing, which is outlined in Supplementary Fig. 2. For the first batch of 6,057 movie stacks, the two-dimensional (2D) class averages calculated from manually picked particles were used as the templates for particle autopicking. Then, the autopicked particles were subjected to several rounds of reference-free 2D classification. 2D class averages of the top views of the complex revealed the presence of 44 centrosymmetric dots along the external rim, suggesting a potential 44-mer form of oligomerization. Particles from good classes were combined and subjected to generate the initial model. After several rounds of 3D classification, the resulting classes without apparent structural deformation in each round were joined. Any duplicate particles in the combined data were removed to make the most of the data, as the particles tended to deform easily. These “classification-join-removing” cycles aimed to enrich the good particles as much as possible. Each round of 3D classification was carried out with C1 symmetry imposed. In the final round, the top view of the complex revealed a 44-mer form of oligomerization, which was consistent with the 2D class averages shown in Supplementary Fig. 3c.

With this prior knowledge, the second batch of 13,420 movie stacks was processed using a similar approach, and 3D classifications were performed with C11 symmetry imposed at the beginning. The most apparent deformation of the particles was observed at the bottom rim of the dome, where they changed from circular to elliptical shapes with varying eccentricities. By imposing high rotational symmetry during 3D classification, classes enriched with elliptical-shaped particles were characterized by features that were easier to observe, such as broken features and incorrect sizes of the bottom rim. Consequently, it facilitated the identification of the most homogeneous classes and the removal of deformed particles.

Alignment of particle images with the same chirality during 3D reconstruction is another challenge. When images with both the correct and inverse chirality coexist in the same map, the resulting resolution is low. To address this issue, a good reference map with a definite chirality was required. Based on the medium-quality C11-symmetric 3D density map, a preliminary 44-mer model was manually constructed and then converted to a map. The artificial map was used as the reference to refine the reconstructed map, resulting in a definite chirality and significant improvement in resolution. A final set of particles, containing 25,977 particles from two datasets, were pooled together and subjected to 3D refinement, resulting in a 3.57 Å (Gold-standard FSC 0.143) map with C22 symmetry imposed, a 6.32 Å (Gold-standard FSC 0.143) map with C1 symmetry imposed.

### Model building and refinement

Flotillin-1 and flotillin-2 were modeled de novo using Coot ^68^, partially based on the AlphaFold2 ^69,70^ predicted monomer structures.

A poly-alanine model was built using Coot, and sequence substitution was performed using bulky side-chain residues as makers. Models of SPFH domain were generated by performing rigid body fitting of the AlphaFold2 predicted structures into the density map, followed by manual adjustments using Coot.

The 1/11 map, containing two flotillin-1 and two flotillin-2 subunits, was used for model refinement using real-space refinement (phenix.real_space_refine) in Phenix^71^. Model validation (Table S1) was calculated by MolProbity^72^. Figure preparation and structure analysis were performed with UCSF Chimera^73^.

### Cryo-ET sample preparation, cryo-FIB milling, tomographic tilt series collection

The resuspended Jurkat stable cell line described above were loaded onto Quantifoil grids (R2/2, Au 200-mesh grid, Quantifoil Micro Tools, Germany), which were freshly glow discharged with a plasma cleaner before use. Excess liquid was blotted away from the back, and grids were plunge-vitrified with the Thermo Fisher Vitrobot Mark IV system. The vitrified samples were stored in liquid nitrogen until milling.

To prepare lamellae at the region of interest, cryo-FIB milling was performed following a previous protocol ^74^. Briefly, cryo-fluorescence microscopy was performed with Thunder Imager EM Cryo CLEM (Leica) equipped with a 50×/0.9 NA objective to identify regions of interest. Then samples were loaded into the Thermo Fisher Aquilos 2 Cryo-FIB system using a cryo-transfer rod. To reduce the accumulation of electrons on samples and provide extra support while milling, sputtering was first performed on the samples with a 30 mA current for 15 s followed by organometallic platinum (Pt) deposition via GIS for 20 s. The fluorescence image was overlayed with the SEM images to locate the region of interest for milling. A 500 pA current was used for rough milling around the target region, and then the current was reduced gradually to 10 pA for fine milling, until the thickness of the lamellae reached ∼200 nm.

The milled lamellae were loaded into to a Thermo Scientific Titan Krios TEM equipped with a Falcon 4i camera and a Thermo Scientific Selectris X energy filter. Automatic tomographic tilt series acquisition was performed using Tomography 5.12.0 (ThermoFisher Scientific) dose symmetrically ^75^ from -56° to +56° (with respect to the pre-tilt angle) in 2° increments. 80 sets of images were acquired at a magnification of 42,000 × (pixel size 3.00 Å) using dose fractionation mode. The defocus range was set from -3.0 to -5.0 μm. The total electron dose per tilt series was limited to 120 e^-^/Å^2^.

### Tomogram reconstruction, annotation and rendering

We used the TOMOMAN script for batch tomogram reconstruction^76^. The collected frames were motion-corrected using MotionCor2 software^65^. Subsequent reconstruction was performed using IMOD software, where the tilt series were aligned by patch tracking, and then reconstructed using the weighted back projection method^77^. For visualization, the tomograms were downscaled using a binning factor of 4 and filtered to improve the contrast^78^.

Membrane segmentation was performed automatically using a CNN based MemBrain-seg software^79^, followed by manual polishing with Amira 2022.2 (ThermoFisher Scientific). Actin in the tomograms was localized by the fiber tracking module in Amira (ThermoFisher Scientific), where a reference cylinder with a radius of 3.5 nm and a length of 40 nm was used for template matching. The results were rendered using UCSF Chimera^73^.

### Template matching and subtomogram averaging (STA)

We used the MATLAB (Mathworks) TOM toolbox^80^ to perform general preprocessing of the tomograms. Then the SPA structure of the flotillin complex obtained in our work was low-pass filtered as an initial template. Template matching was performed in the tomograms using the template above through PyTom^81^. In total, 218 flotillin complexes were selected from 20 tomograms. The subtomograms with a box size of 176 pixels^3^ (1× binned) were extracted using Warp^78^, then aligned in RELION v3.1^82^, where C11 symmetry was applied. The model of flotillin complex was fitted into the STA map using rigid body docking within UCSF Chimera^73^. For visulization, the averaged structures were mapped back to the tomograms in their original locations and orientations, using the information from template matching and subtomogram averaging.

### Comparative lipidomics analysis

Plasma membrane fractions were isolated following a previous publication^83^. In brief, HEK293F cells (10 mL) were pelleted and resuspended in 2 mL phosphate-buffered saline (PBS). The cells were lysed by 50 strokes in a Dounce homogenizer. Nuclei and cell debris were removed by centrifugation at 1000 × g for 10 min. The resulting supernatant (2 mL) was layered onto 8 mL of 30% Percoll in PBS and centrifuged at 84000 × g for 30 min. The plasma membrane fraction was collected from the membrane band approximately halfway down the gradient.

For lipid extraction, MTBE/methanol extraction method was used. Briefly, purified flotillin protein sample or plasma membrane fraction containing 0.5–2 µg protein was diluted with Buffer E or PBS to a final volume of 100 μL, respectively. The sample was mixed with 300 μL methanol and vortexed. Subsequently, 1 mL of MTBE containing 0.3 µg of each internal standard (LPC (16:0)-D49, LPE (17:0)-D5, PC (16:0-18:1)-D31, PE (16:0-18:1)-D31, PS (16:0-18:1)-D31 from Avanti Research, and TAG (15:0-18:1-15:0)-D7, DAG (15:0-18:1)-D7, Cer (d18:1-22:0)-D7, SM (18:1-18:1)-D9, Cholesterol-D7 from Alta Scientific) was added and vortexed again, followed by adding 250 µL of H₂O to facilitate phase separation. The vortexing and resting process was repeated three times to ensure thorough extraction. The mixture was centrifuged at 8,000 rpm for 15 minutes. The upper organic phase was carefully transferred into glass vials and dried under an argon flow in a fume hood. Aspirating precipitated protein in the interphase and the lower phase should be avoided during collection of the upper phase. Ensure the argon flow is gentle and avoid applying heat during the drying process. All steps above were performed at room temperature. The dried samples were stored at -80 °C for subsequent LC-MS/MS analysis.

For quantitative analysis of general lipids, LC-MS/MS was conducted using an Orbitrap OE240 mass spectrometer coupled with an Ultra-High Performance Liquid Chromatography (UHPLC) system (Thermo Fisher Scientific, USA). The liquid chromatography (LC) separation was performed on a Cortecs C18 column (2.1 × 100 mm, Waters, USA). The LC gradient utilized 60% acetonitrile with 5 mM ammonium acetate as mobile phase A and isopropanol/acetonitrile (v/v 9:1) as mobile phase B. Data-dependent MS/MS acquisition mode was employed, with an MS resolution of 60,000 and an MS/MS resolution of 15,000. The top 10 most intense precursor ions were selected for fragmentation. The mass range for MS acquisition was set to 240–2000 m/z in positive ion mode. Lipid identification was performed using LipidSearch 4.2 software (Thermo Fisher Scientific, USA). Lipids were identified by matching MS/MS spectra with the database in the software. Results with low MS/MS matching scores or inconsistent chromatographic behavior were manually reviewed and excluded to ensure the reliability of lipid identification.

For analysis of cholesterol, LC-MS/MS was conducted using a Q-Exactive HFX mass spectrometer coupled with a UHPLC system (Thermo Fisher Scientific, USA). A 2 µL aliquot of the supernatant was loaded to a Kinetex® Biphenyl column (2.1 × 150 mm, 2.6 µm, Phenomenex). Samples were eluted using acetonitrile containing 0.1% formic acid as the eluent, with a gradient from 80% to 99% within 6.5 min. The stationary phase was aqueous with 0.1% formic acid. Data acquisition was performed in positive ion mode with a mass range of 300–500 m/z using data-dependent MS/MS. Full scan and fragment spectra were collected at mass resolutions of 60,000 and 15,000, respectively. The source parameters were set as follows: spray voltage, 3,200 V; capillary temperature, 320 °C; heater temperature, 300 °C; sheath gas flow rate, 35 Arb; auxiliary gas flow rate, 10 Arb. Data analysis was performed using Xcalibur 4.4 software (Thermo Fisher Scientific, USA). Cholesterol assignment was confirmed using a chemical standard based on retention time.

The molar amount of each lipid species was determined using corresponding internal lipid standards. For lipids without specific internal standards, calculations were based on similar lipid species: PI and PG were estimated using the PS standard, and SPH was estimated using the Cer standard. For every sample, the molar amounts of all lipid species, including cholesterol, were summed, and each lipid species was normalized to its mole percentage of the total lipid content.

### Statistical tests

Statistical analyses were conducted using GraphPad Prism software. To evaluate the differences between two independent groups (purified flotillin protein sample and plasma membrane fraction sample) (Fig. 5d), unpaired two-sample t-tests were applied with Welch’s correction, which does not assume equal variances between groups. Each kind of lipid species was treated as an independent comparison, and multiple t-tests were performed without assuming consistent standard deviations across groups. All statistical tests were two-tailed, assumed a Gaussian distribution, and parametric methods were applied.

## Statistics and Reproducibility

All experiments were performed with appropriate replicates to ensure statistical validity. Data presented in Fig. 2i, Fig. 4a, b and Supplementary Fig. 7b are based on three independent experiments.

## Data availability

The cryo-EM map generated in this study have been deposited in the Electron Microscopy Data Bank (EMDB) under accession code EMD-62785 [https://www.ebi.ac.uk/pdbe/entry/emdb/EMD-62785] (structure of the flotillin complex); EMD-67802 [https://www.ebi.ac.uk/pdbe/entry/emdb/EMD-67802] (structure of the flotillin complex in situ);. The atomic coordinate generated in this study have been deposited in the Protein Data Bank (PDB) under accession code 9L3G [https://doi.org/10.2210/pdb9L3G/pdb] (structure of the flotillin complex). The lipidomics data generated in this study are provided in the Supplementary Data file. The source data underlying Figure 2i, and Supplementary Figures 1a, 2a, 7b and 8d are provided as a Source Data file.

## Acknowledgements

We thank Dr. Chengying Ma and Dr. Ningning Li for advice in cryo-EM data processing, Xing Wang, Wenjing Du, Dr. Changdong Qin and Dr. Wenhong Jiang for aiding in cryo-ET experiments, and all members of the Gao Lab and Guo Lab for insightful discussions. We thank the Core Facilities at the School of Life Sciences, Peking University for help with negative staining EM and confocal microscopy; the Cryo-EM Platform and the Electron Microscopy Laboratory of Peking University for help with data collection; the High-performance Computing Platform of Peking University for help with computation; and the Facility Center of Metabolomics and Lipidomics, National Center for Protein Sciences, Tsinghua University for help with lipidomics analysis; and the National Center for Protein Sciences at Peking University for assistance in key experiments. This work was supported by National Natural Science Foundation of China (Grant # 92354306), and by the Ministry of Science and Technology of China and Changping Laboratory. N. G. is supported in part by the Frontier Innovation Fund of Peking University Chengdu Academy for Advanced Interdisciplinary Biotechnologies. Q.G. is supported by the Beijing Natural Science Foundation (Grant # JQ24031).

## Author contributions

N. G. and Q. G. designed and supervised the experiments; M. L. purified the sample with assistance from L. M. and J. H. and conducted all other experiments. Y. Q. and M. L. performed cryo-ET data analysis. X. L. and L. Y. assisted with the work. M. L. prepared figures and wrote the manuscript. N. G. and Q. G. revised the manuscript. All the authors approved the manuscript.

