## Supplementary Information for "Molecular mechanisms of flotillin complexes in organizing membrane microdomains"

### Supplementary Figure 1

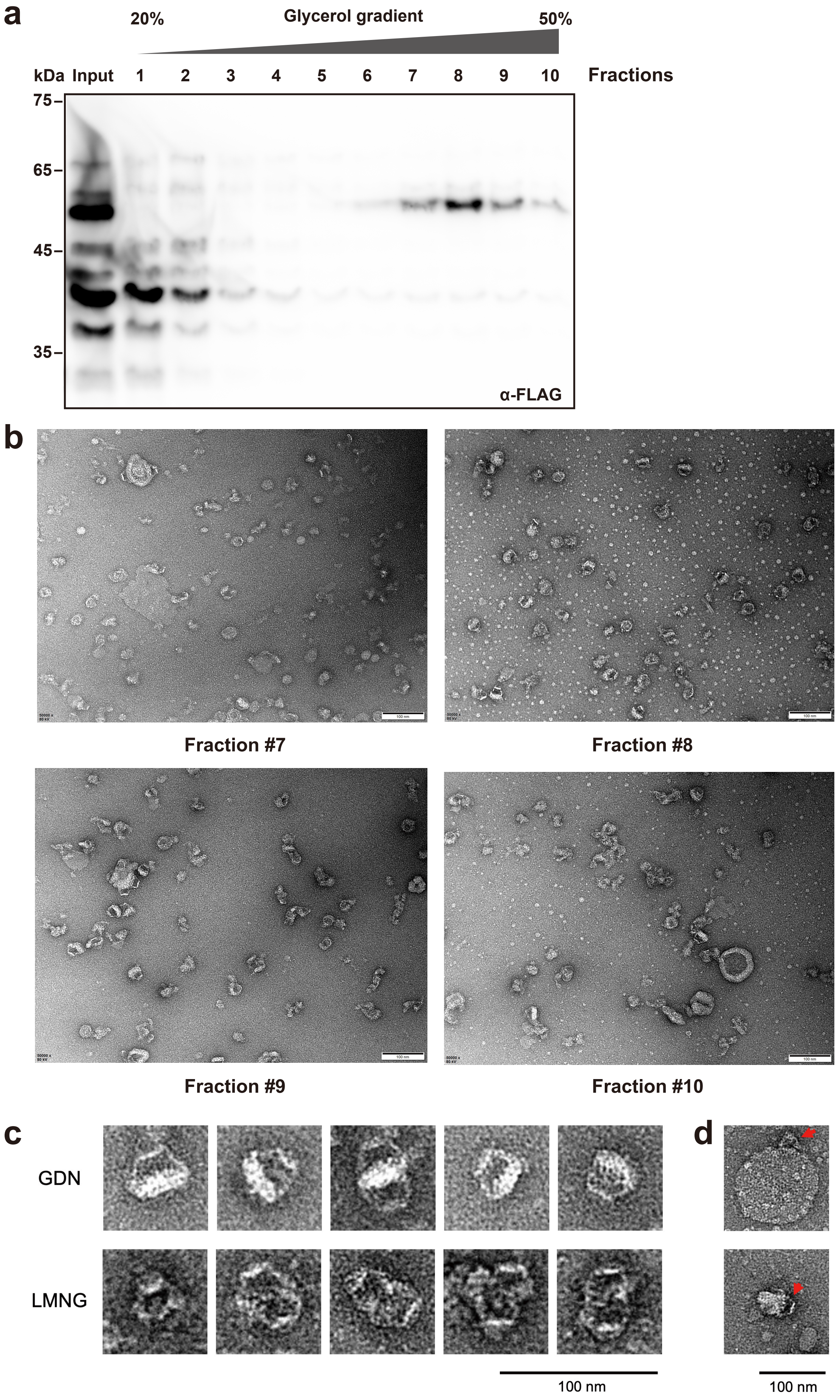

**Supplementary Figure 1. Purification of the flotillin complex.**

**a,** Western blot (WB) analysis shows the distribution of the flotillin complex in the glycerol gradient. Flotillin-1 contains a C-terminal FLAG tag, and flotillin-2 has no tag. Source data are provided as a Source Data file.

**b,** The negative staining electron microscopy (nsEM) analysis of the samples from tube 7 to tube 10 in **a**. Almost trapezoidal particles are the side view of the flotillin complex. Some of them attach to the vesicles through the bottom of the trapezoid.

**c,** 2D nsEM images of particles purified with GDN or LMNG detergents, respectively. Under the condition of GDN, particles tend to retain the membrane inside the complex. Under the condition of LMNG, particles tend to dimerize through the membrane associated region, probably due to the absence of the membrane.

**d**, The nsEM images show flotillin complexes attached to vesicles in purified sample.

Scale bars, 100 nm (**b–d**).

### Supplementary Figure 2

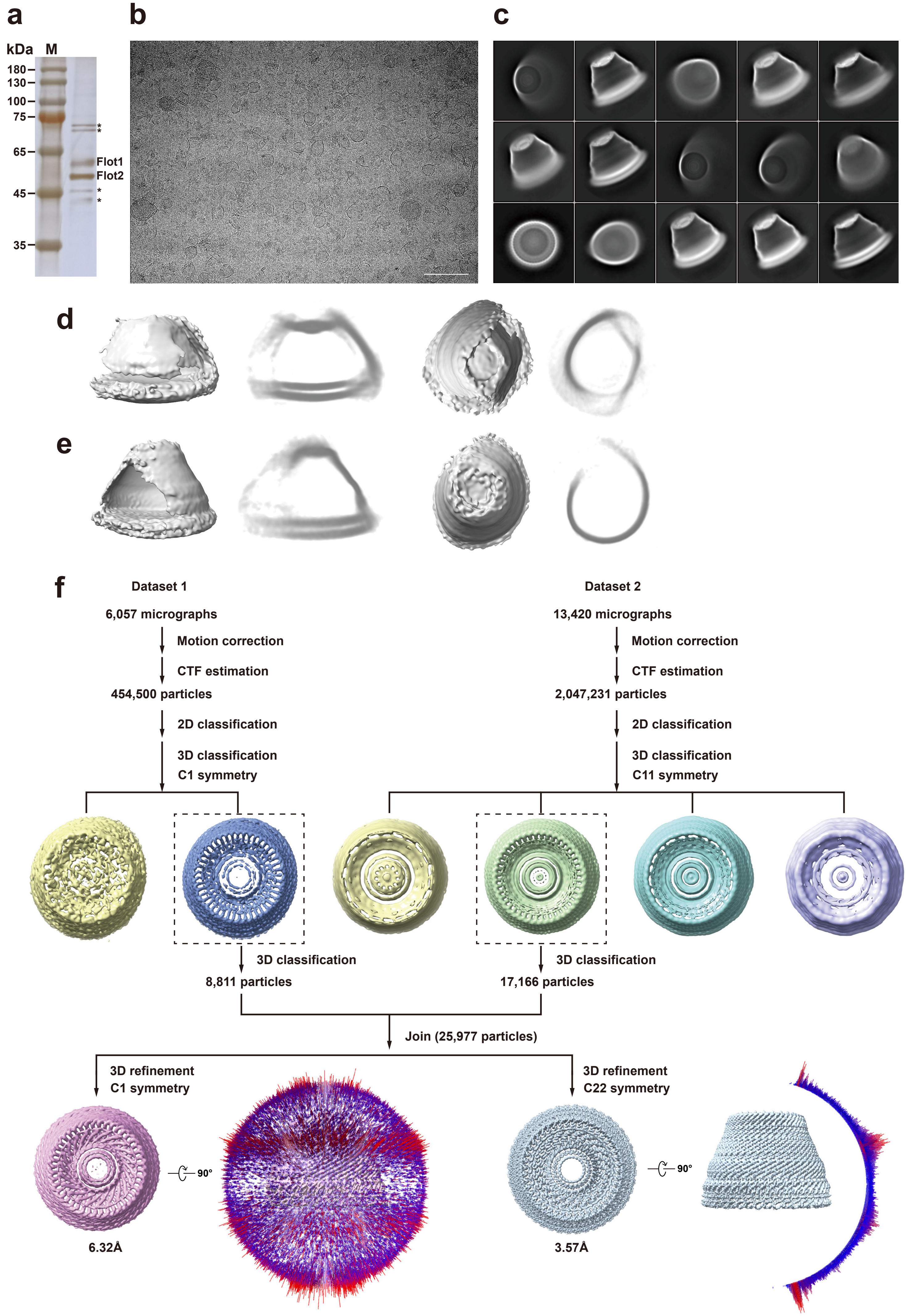

**Supplementary Figure 2. Structural determination of the flotillin complex.**

**a,** Silver stained SDS-PAGE gel of the flotillin complex used for the cryo-EM analysis. Source data are provided as a Source Data file.

**b,** A representative cryo-EM image of the flotillin complex.

**c,** Representative average images of 2D classification. The image at the bottom left is the top view of the flotillin complex, in which feature of 44-mer oligomerization is almost visible.

**d & e,** Side views and top views of deformed maps during cryo-EM SPA 3D classification. The slabs of these maps are shown aside.

**f,** The flow chart of image processing for the two cryo-EM data batches. Particles with apparent feature of 44-mer oligomerization were pooled for high-resolution refinement with C1 and C22 symmetry imposed. Angular distributions of the particles used in the final reconstructions (C1 and C22) are displayed alongside their respective maps.

Scale bars, 100 nm (**b**).

#### Supplementary Figure 3

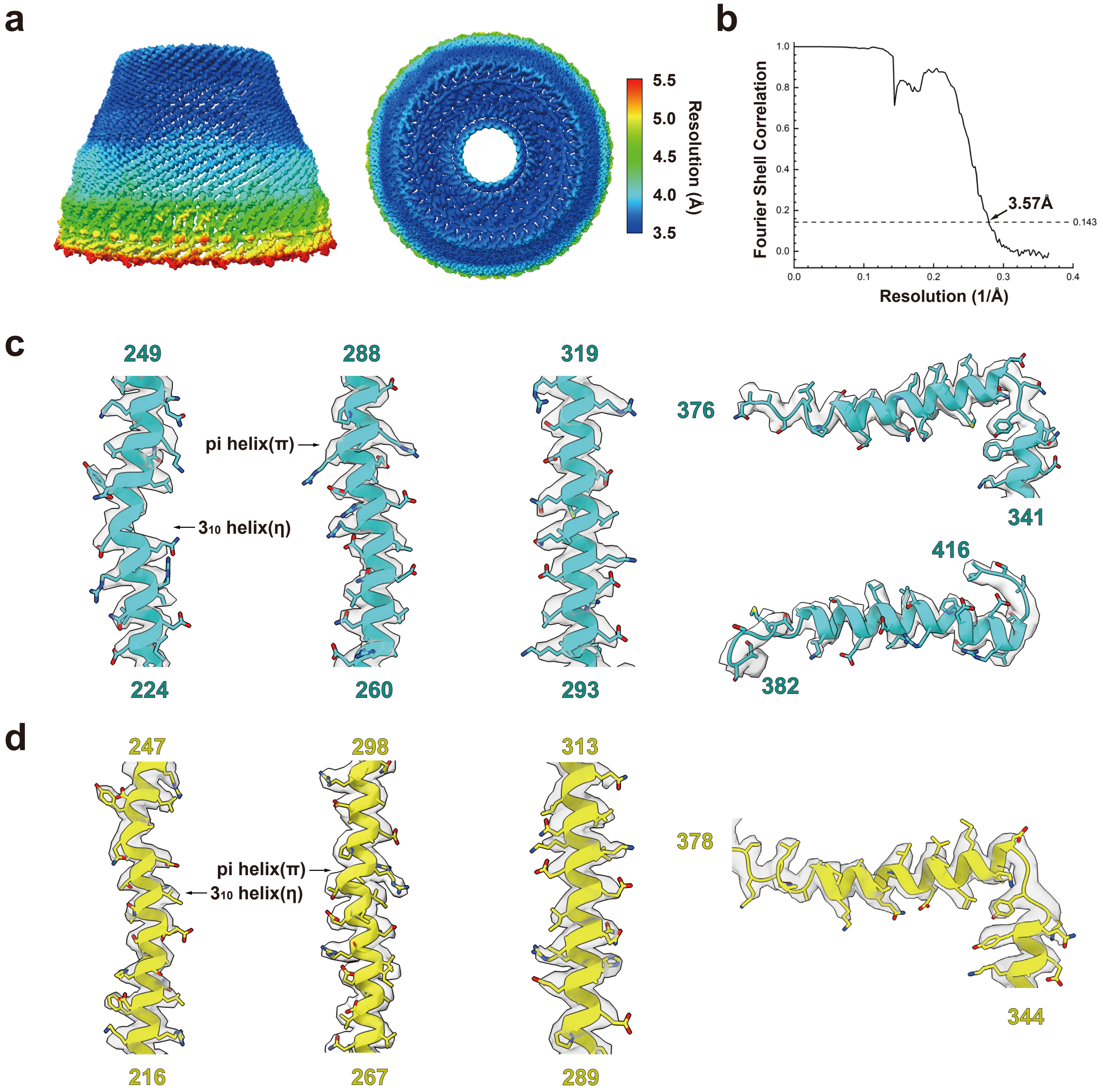

**Supplementary Figure 3. Resolution estimation of the final density map of the flotillin complex.**

**a,** Local resolution estimation of the final cryo-EM map.

**b,** Fourier shell correlation (FSC) curves of the final map. At FSC gold-standard 0.143 cutoff, the estimated resolution is 3.57 Å.

**c & d,** Local densities of selected regions of flotillin-1 (**c**) and flotillin-2 (**d**), superimposed with respective atomic models. Pi helices ($\pi$) and $3_{10}$ helices ($\eta$) are labeled.

### Supplementary Figure 4

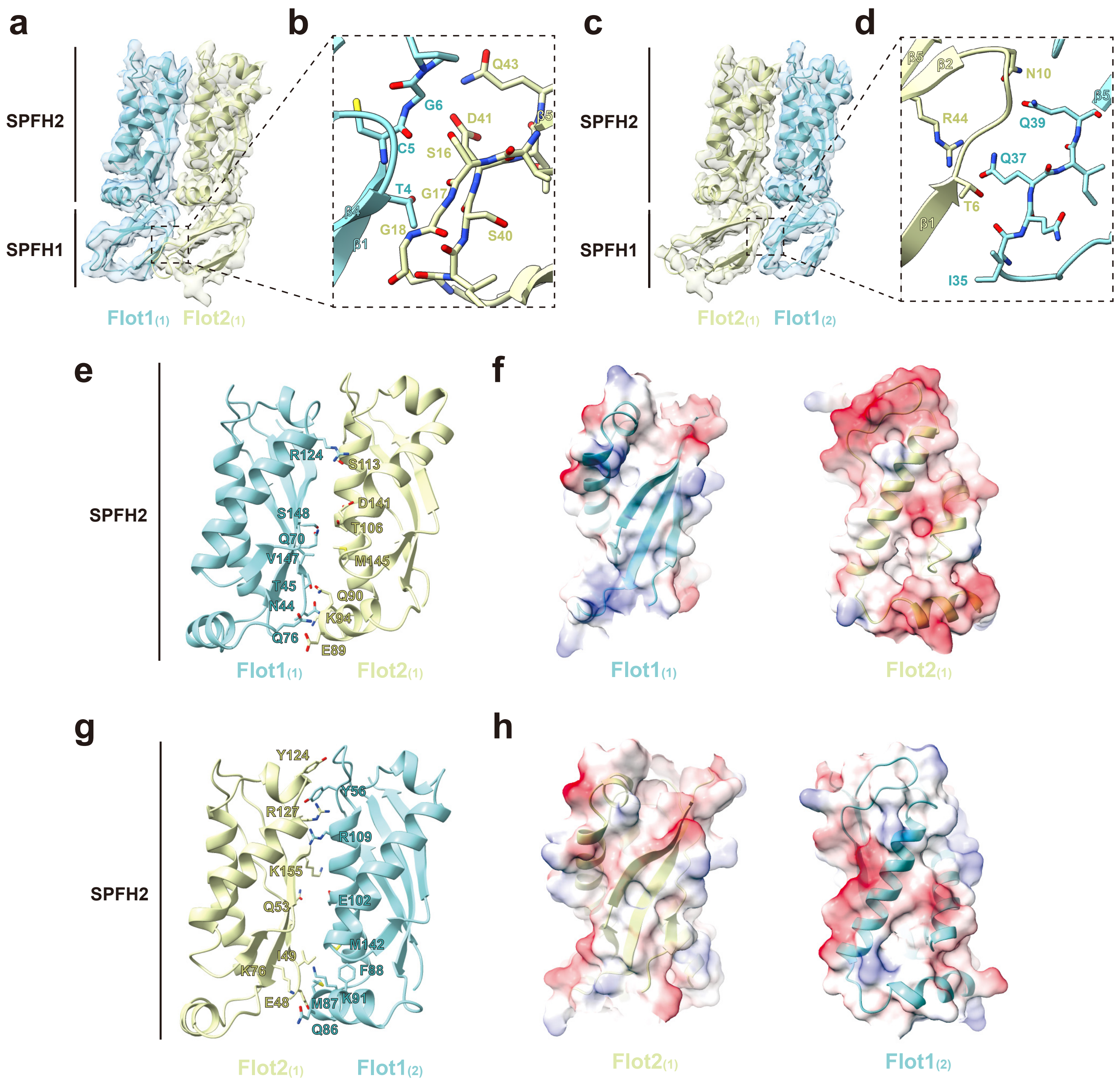

**Supplementary Figure 4. Interactions between SPFH domains of flotillin-1 and flotillin-2.**

**a & c,** Domain interface between $F1_{(1)}$ and $F2_{(1)}$ (**a**), between $F2_{(1)}$ and $F1_{(2)}$ (**c**) in the SPFH1 region. The cryo-EM density maps (transparent surfaces) are shown with corresponding backbone models (solid ribbons) derived from reliable tracing of the SPFH1 and SPFH2 domains.

**b & d,** Close-up views demonstrate that intermolecular interactions between flotillin-1 and flotillin-2 SPFH1 domains are mediated by loop regions at two distinct interfaces, $F1_{(1)}$:$F2_{(1)}$ interface (**b**) and $F2_{(1)}$:$F1_{(2)}$ interface (**d**).

**e,** The $F1_{(1)}$-$F2_{(1)}$ interface is shown in ribbon representation, with interacting residues highlighted in stick models.

**f,** Electrostatic surface potential of the interface between the SPFH2 domains of $F1_{(1)}$ and $F2_{(1)}$.

**g & h,** Same as **e** & **f**, for the $F2_{(1)}$-$F1_{(2)}$ interfaces.

### Supplementary Figure 5

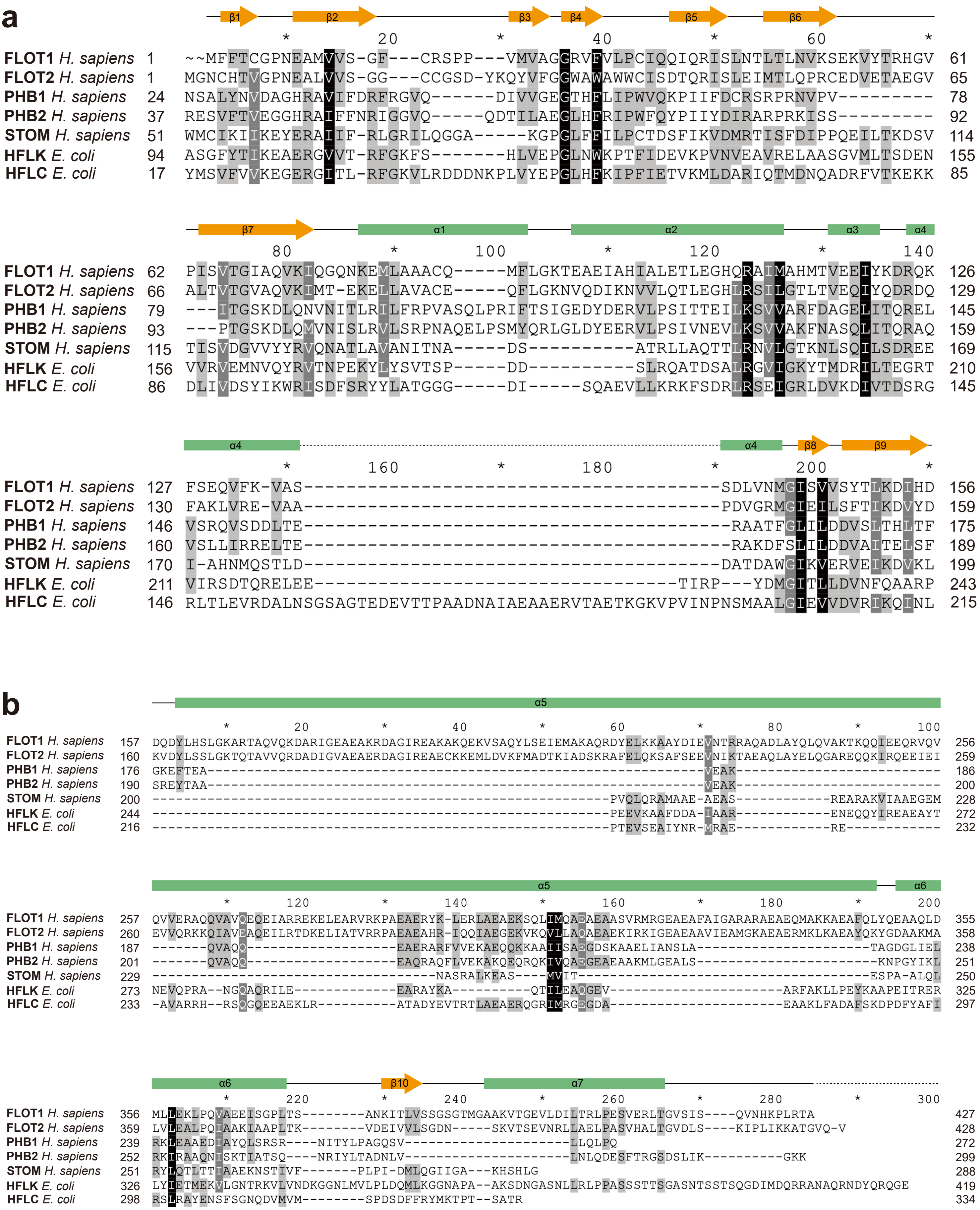

**Supplementary Figure 5. Multiple sequence alignment of representative SPFH family members.**

**a & b**, Sequence alignment of the SPFH domains (**a**), CC1, CC2 and CTD domains (**b**) from a few SPFH family members, including *H. sapiens* flotillin-1 and -2, prohibitin-1 and -2, stomatin, *E. coli* HflC and HflK. The secondary structure motifs of flotillin-1 were labelled. Protein sequences in the SPFH domains (**a**), CC2 (α6) and CTD domains (β10 and α7) (**b**) are more conserved, whereas that in the CC1 domains are less conserved. The length of the CC1 subdomains (α5) in flotillin-1 and flotillin-2 are same and much longer than those in the other SPFH family members.

### Supplementary Figure 6

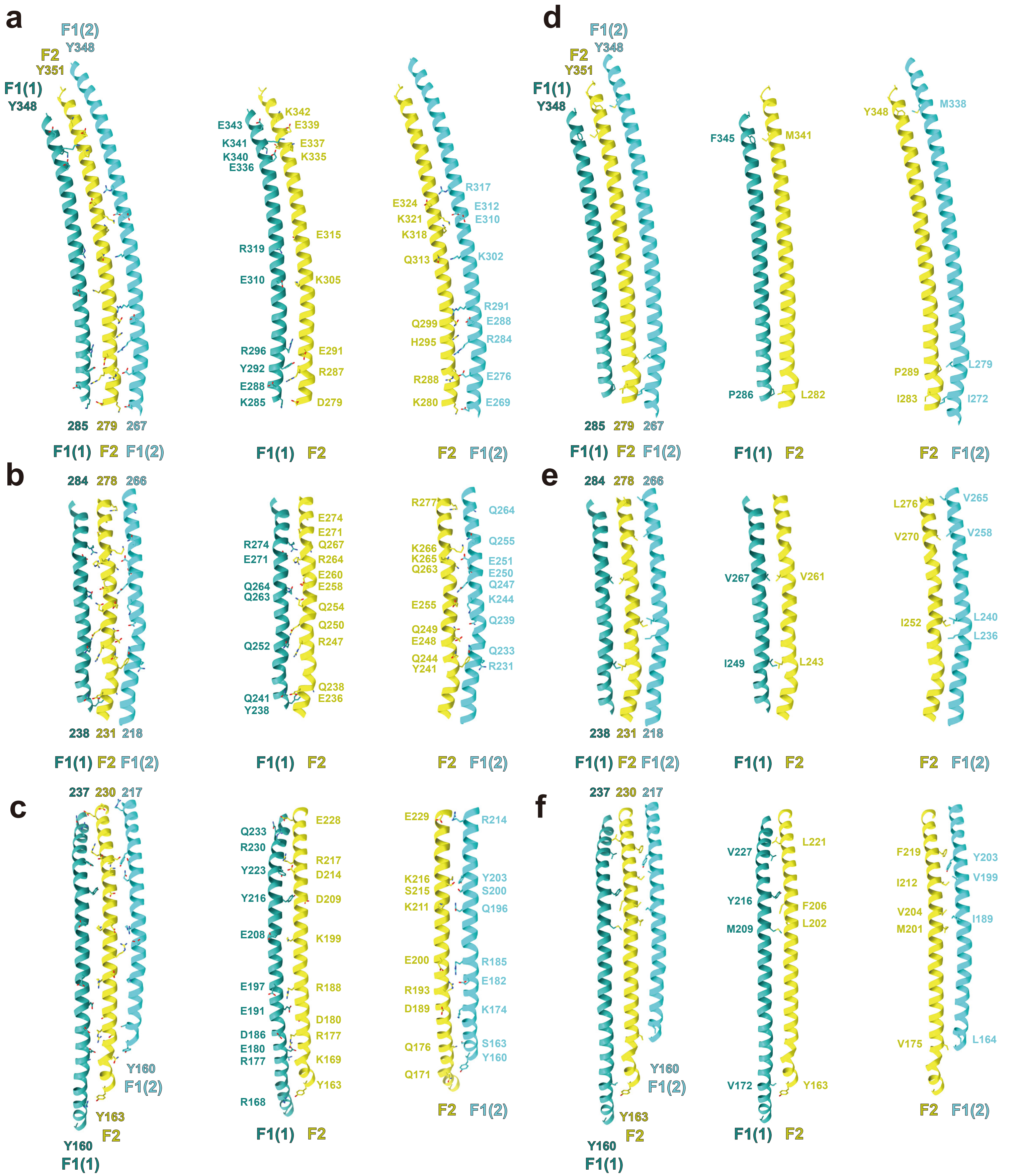

**Supplementary Figure 6. Interactions between CC1 subdomains of flotillin-1 and flotillin-2.**

**a–c,** Hydrophilic interactions between CC1 domains of flotillin-1 and flotillin-2, with participating residues highlighted in stick models, for the $F1_{(1)}$-F2, F2-$F1_{(2)}$ interfaces. Due the large range of bending of the CC1 subdomains, they were cut into three segments along the perpendicular line of the helices, facilitating to show in detail.

**d–f,** Same as (**a–c**), but for the hydrophobic interactions.

### Supplementary Figure 7

**a**

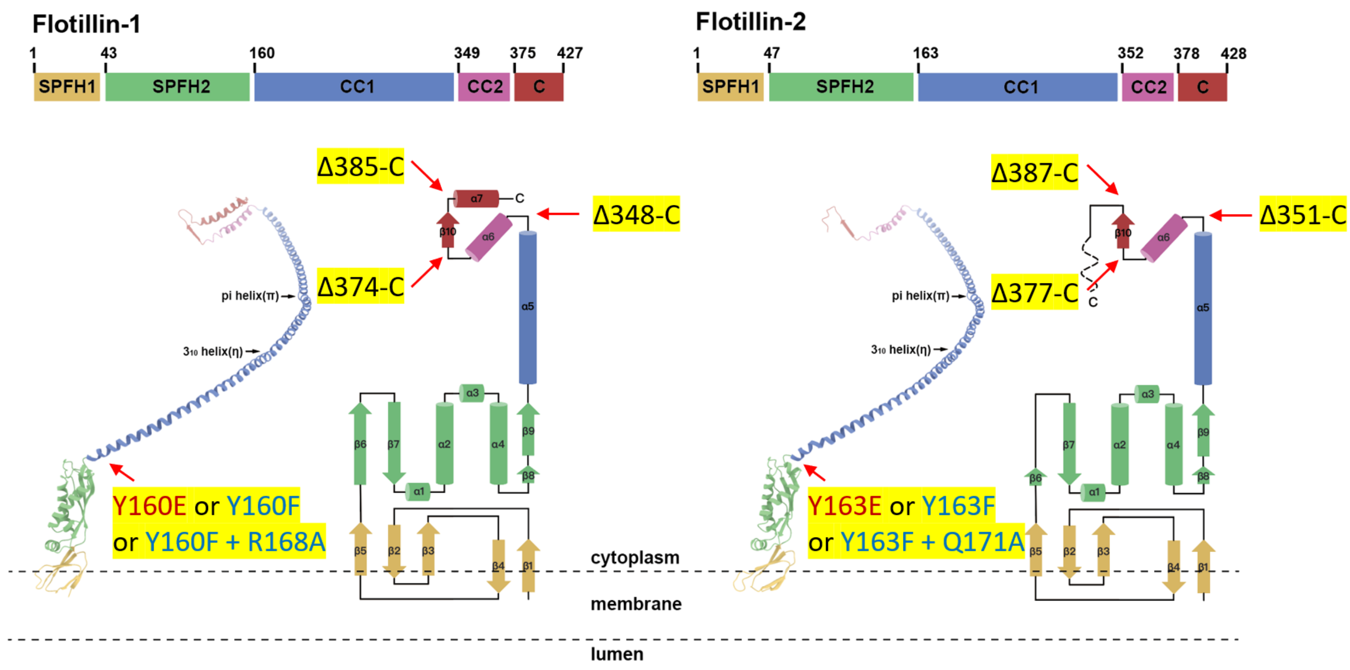


**b**

| | F1-Flag | F2-Strep | kDa α-Flag (20% → 50%) | kDa α-Strep (20% → 50%) |
|---|---|---|---|---|
| i | WT | - | 1 2 3 4 5 6 7 8 9 10 11<br>65 - 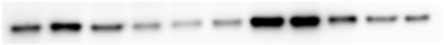 | N/A |
| ii | - | WT | N/A | 1 2 3 4 5 6 7 8 9 10 11<br>65 - 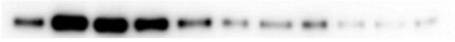 |
| iii | Y160E | - | 1 2 3 4 5 6 7 8 9 10 11<br>65 - 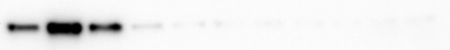 | N/A |
| iv | - | Y163E | N/A | 1 2 3 4 5 6 7 8 9 10 11<br>65 - 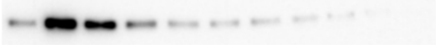 |
| v | Y160F R168A | Y163F Q171A | 1 2 3 4 5 6 7 8 9 10 11<br>65 - 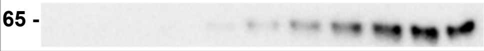 | 1 2 3 4 5 6 7 8 9 10 11<br>65 - 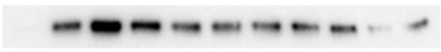 |
| vi | Δ385-C | Δ387-C | 65 - 1 2 3 4 5 6 7 8 9 10 11<br>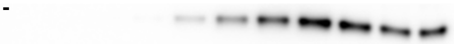 | 65 - 1 2 3 4 5 6 7 8 9 10 11<br>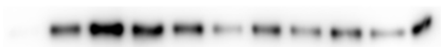 |
| vii | Δ374-C | Δ377-C | 65 - 1 2 3 4 5 6 7 8 9 10 11<br>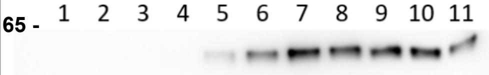 | 65 - 1 2 3 4 5 6 7 8 9 10 11<br>45 - 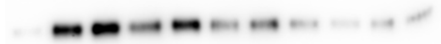 |
| viii | Δ348-C | Δ351-C | 65 - 1 2 3 4 5 6 7 8 9 10 11<br>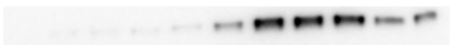 | 1 2 3 4 5 6 7 8 9 10 11<br>45 - 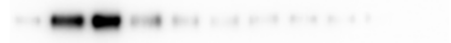 |

**Supplementary Figure 7. Analysis of dome structure formation in wild-type and mutant flotillins.**

**a,** Schematic diagrams show the mutations and truncations of flotillin-1 and flotillin-2, respectively.

**b,** Glycerol gradient fractionation analysis of crude membrane extracts expressing WT or mutant flotillin constructs (expressed individually or in combination; experimental setup identical to Figure 2i). In this series of experiments, flotillin-1 was C-terminally Flag-tagged while flotillin-2 contained a C-terminal Strep tag. Source data are provided as a Source Data file.

### Supplementary Figure 8

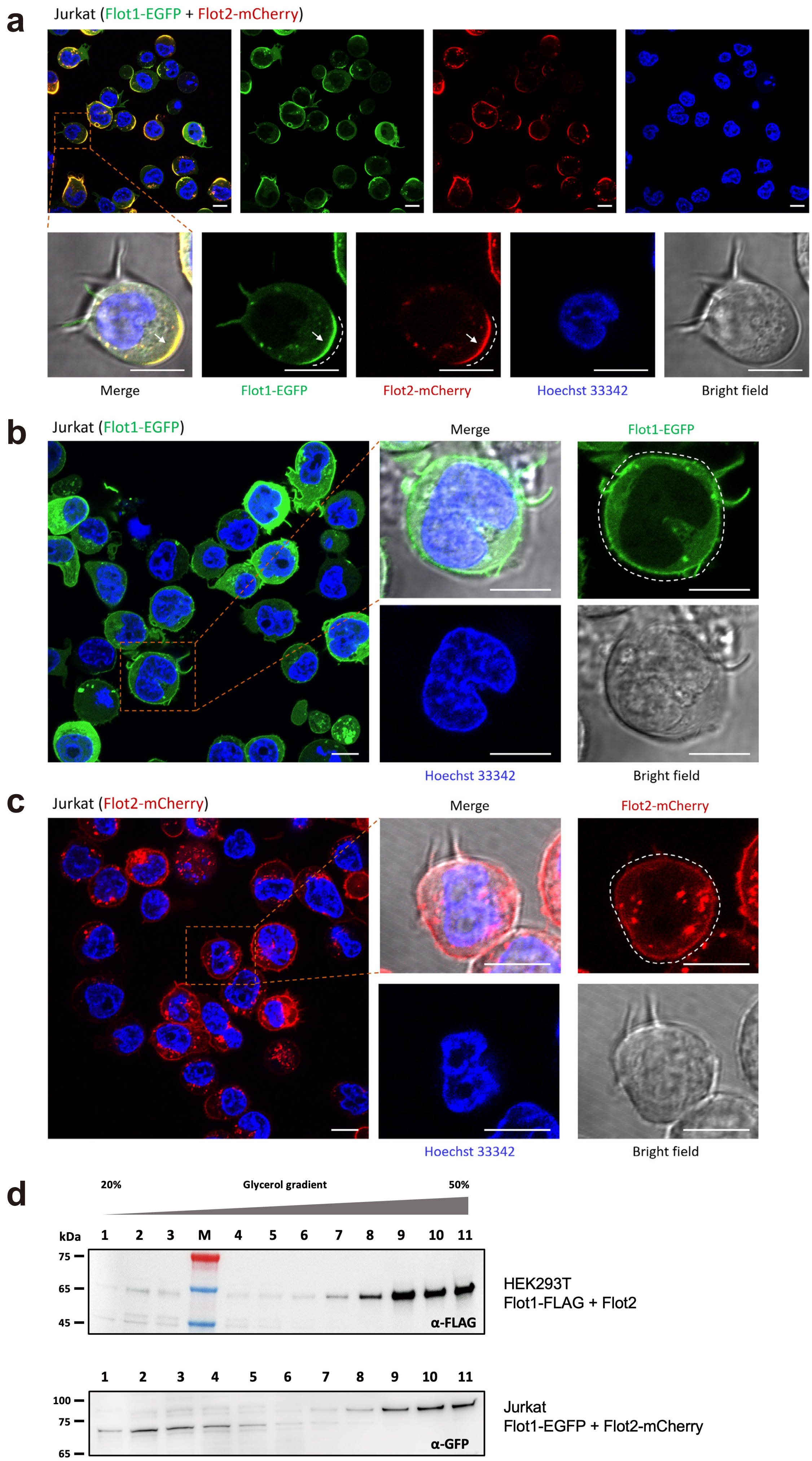

**Supplementary Figure 8. Fluorescent protein tags have no effect on flotillins oligomerization.**

**a–c,** Subcellular distribution of flotillin-1 and flotillin-2 in Jurkat cells.

**a**, Co-expressed Flot1-EGFP (green) and Flot2-mCherry (red) show polarized accumulation at uropods, with dashed arcs and arrows highlighting the enrichment pattern (enlarged regions).

**b & c**, When expressed individually, Flot1-EGFP (**b**) and Flot2-mCherry (**c**) display uniform plasma membrane distribution, as indicated by dashed outlines (enlarged regions). Nuclei were stained with Hoechst 33342 (blue).

**d,** The density gradient centrifugation analysis was done as described in Extended Data Fig. 7b for the F1-EGFP and F2-mCherry pair. They oligomerized as the WT flotillins did. Source data are provided as a Source Data file.

Scale bars, 10 µm (**a–c**).

#### Supplementary Figure 9

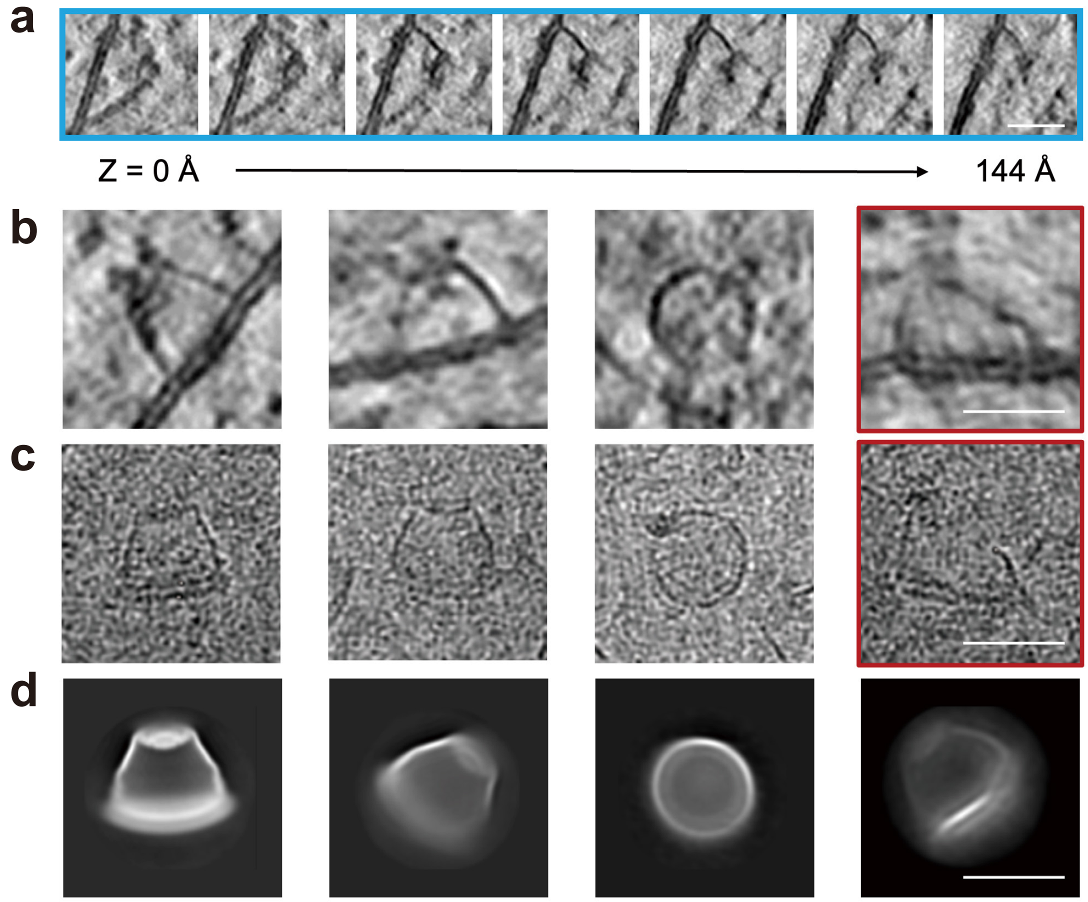

**Supplementary Figure 9. Structural plasticity of the flotillin complex.**

**a,** A series of tomographic slices zoomed in on an individual flotillin complex detected in one tomogram. One side of the flotillin complex is deformed and the density in this region is attenuated.

**b–d,** Representative particles from slices of tomograms (**b**), cryo-EM micrographs (**c**) and 2D averages during cryo-EM SPA (**d**). Red boxes indicate similar particles with significant degree of deformation.

Tomographic slices are 1.2 nm thick. Scale bars, 30 nm (**a**–**d**).

### Supplementary Figure 10

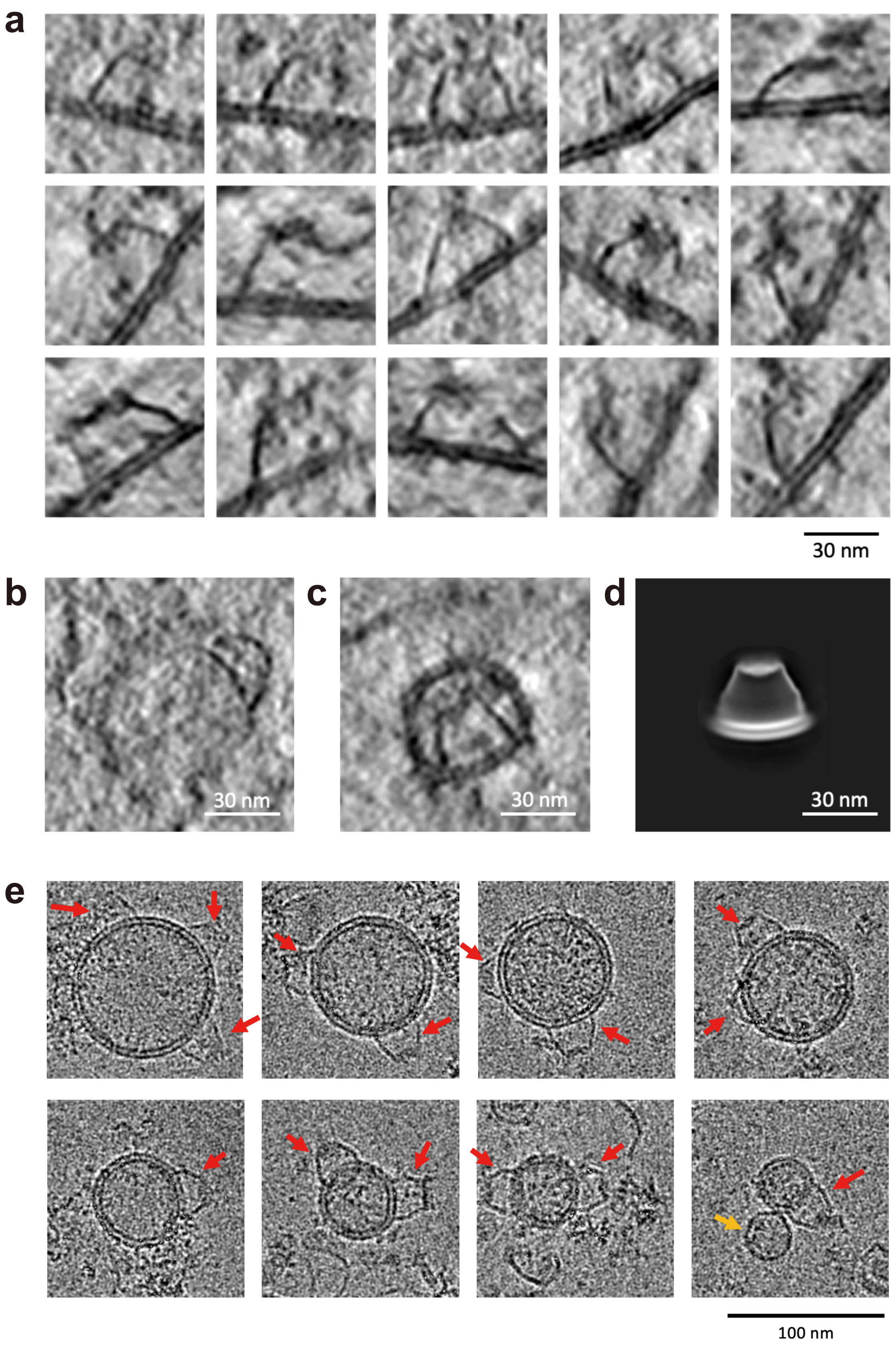

**Supplementary Figure 10. The curvature of the membrane covered by the flotillin complex.**

**a,** Tomographic slices zoomed in on the flotillin complex show nearly flat membranes covered by the flotillin complex in situ.

**b,** The flotillin complex covered the convex membrane of an endosome found in the tomogram described in Fig. 5b.

**c,** A flotillin complex on a highly concave membrane surface inside a tiny vesicle, which was inside an apoptosis cell and undergoing degradation in endolysosomes according to the cellular context in situ.

**d,** The 2D averaged image during cryo-EM SPA shows that membranes retained inside the purified the flotillin complex tend to form a concave shape.

**e,** The cryo-EM images show that the flotillin complex covered the convex of vesicles in purified sample. However, because of the affinity purification step, only vesicles with the flotillin complex inserted in outer leaflet were enriched in final product.

Tomographic slices are 1.2 nm thick (**a**–**c**). Scale bars, 30 nm (**a**–**d**) and 100 nm (**e**).

### Supplementary Figure 11

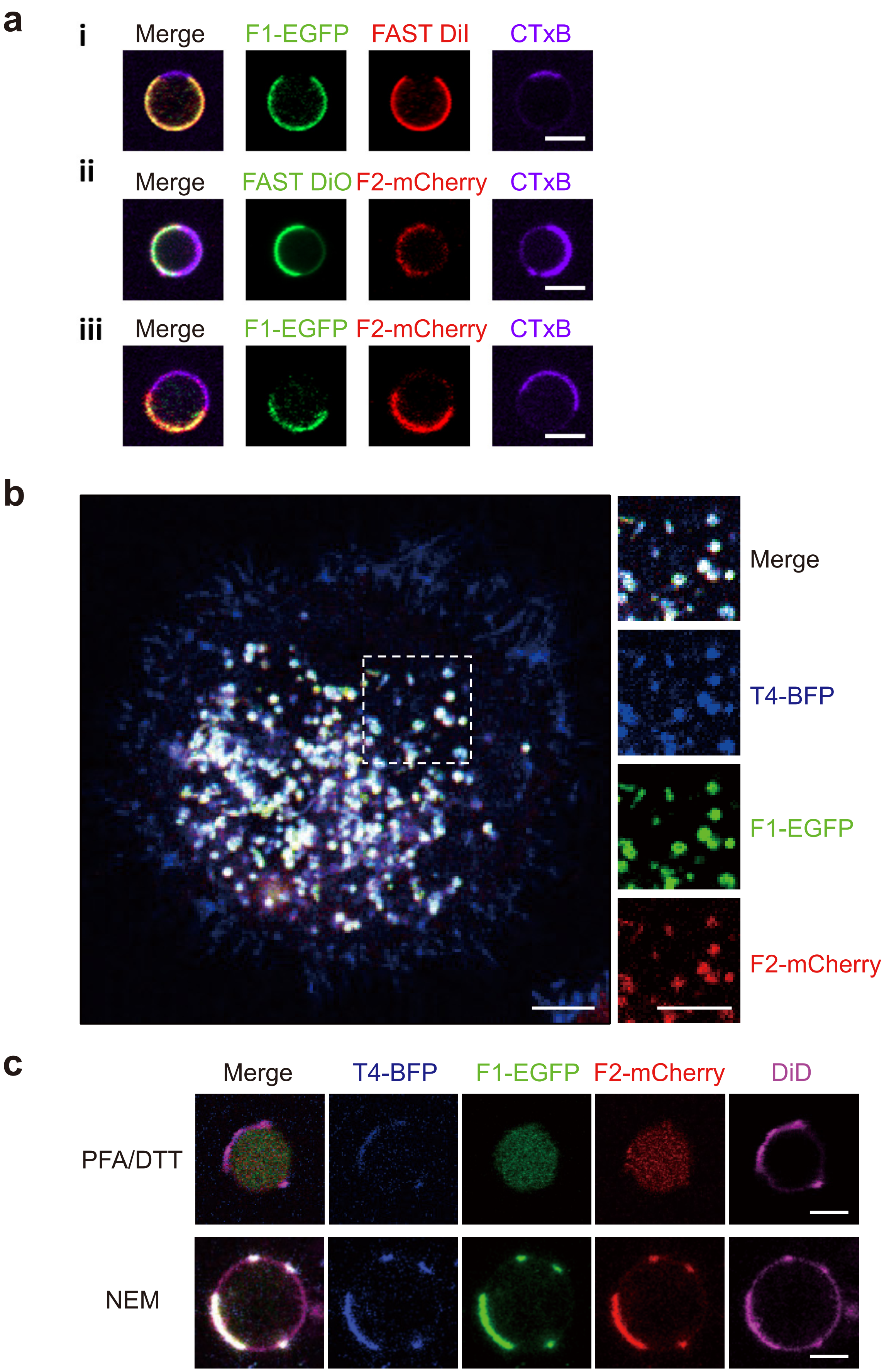

**Supplementary Figure 11. Colocalization of flotillins and tetraspanin-4 in intracellular vesicles and their phase partitioning in GPMVs.**

**a**, GPMVs were labeled with different dyes following isolation from the Jurkat cells expressing Flot1-EGFP (i), Flot2-mCherry (ii), both Flot1-EGFP and Flot2-mCherry (iii). Phase separation into the Lo and Ld phases was induced by cooling, with FAST DiI and FAST DiO dye marking the Ld phase, and CTxB (Cholera Toxin Subunit B (Recombinant), Alexa Fluor™ 647, Invitrogen) marking the Lo phase.

**b,** In L929 cells stably expression F1-EGFP, F2-mCherry and tetraspanin-4-BFP, flotillins and tetraspanin-4 colocalize within intracellular vesicles.

**c,** GPMVs were labeled with DiD dye following isolation from the L929 cells described in **b**. Phase separation into the Lo and Ld phases was induced by cooling, with DiD dye marking the Ld phase. Under PFA/DTT treatment, F1-EGFP and F2-mCherry dissociated from the membrane due to DTT-induced depalmitoylation and diffused within the GPMVs, while tetraspanin-4-BFP remained in the membrane and partitioned into the Ld phase. Under NEM treatment, F1-EGFP, F2-mCherry and tetraspanin-4-BFP all partitioned into the Ld phase.

Scale bars, 5 µm (**a**–**c**).

**Supplementary Table 1. Cryo-EM data collection, refinement and validation statistics**

| | Flotillin complex (EMDB-62785) (PDB 9L3G) | | Flotillin complex in situ (EMDB-67802) |
|---|---|---|---|
| **Data collection and processing** | | **Data collection and processing** | |
| Magnification | 64,000× | Magnification | 42,000× |
| Voltage (kV) | 300 | Voltage (kV) | 300 |
| Electron exposure ($e^{–}/Å^2$) | 41 | Electron exposure ($e^{–}/Å^2$) | 120 |
| Defocus range (μm) | -1.6 to -2.2 | Defocus range (μm) | -3 to -5 |
| Pixel size (Å) | 1.37 | Tilt angle range(°) | -56 to +56 |
| Symmetry imposed | C22 | Pretilt (°) | +13 |
| Initial particle images (no.) | 2,501,731 | Tilt step angle(°) | 2 |
| Final particle images (no.) | 25,977 | Pixel size (Å) | 3 |
| Map resolution (Å) | 3.57 | Symmetry imposed | C11 |
| FSC threshold | 0.143 | | |
| Map resolution range (Å) | 3.5–6.0 | Initial subtomograms | 218 |
| | | Final subtomograms | 218 |
| **Refinement** | | Map resolution (Å) | 25.6 |
| Initial model used (PDB code) | Ab initio | FSC threshold | 0.143 |
| Model resolution (Å) | | | |
| FSC threshold | | | |
| Model resolution range (Å) | | | |
| Map sharpening *B* factor ($Å^2$) | -103.3 | | |
| Model composition | | | |
| Non-hydrogen atoms | 138,556 | | |
| Protein residues | 17,886 | | |
| Ligands | 0 | | |
| *B* factors ($Å^2$) | | | |
| Protein | 82.85 | | |
| Ligand | 0 | | |
| R.m.s. deviations | | | |
| Bond lengths (Å) | 0.006 | | |
| Bond angles (°) | 1.094 | | |
| Validation | | | |
| MolProbity score | 1.55 | | |
| Clashscore | 6.11 | | |
| Poor rotamers (%) | 0.02 | | |
| Ramachandran plot | | | |
| Favored (%) | 96.65 | | |
| Allowed (%) | 3.35 | | |
| Disallowed (%) | 0.00 | | |
